# Intramuscular mRNA BNT162b2 vaccine against SARS-CoV-2 induces robust neutralizing salivary IgA

**DOI:** 10.1101/2022.02.17.480851

**Authors:** Miri Stolovich-Rain, Sujata Kumari, Ahuva Friedman, Saveliy Kirillov, Yakov Socol, Maria Billan, Ritesh Ranjan Pal, Peretz Golding, Esther Oiknine-Djian, Salim Sirhan, Michal Bejerano Sagie, Einav Cohen-Kfir, Maya Elgrably-Weiss, Bing Zhou, Miriam Ravins, Yair E Gatt, Kathakali Das, Orly Zelig, Reuven Wiener, Dana G Wolf, Hila Elinav, Jacob Strahilevitz, Dan Padawer, Leah Baraz, Alexander Rouvinski

## Abstract

Intramuscularly administered vaccines stimulate robust serum neutralizing antibodies, yet they are often less competent in eliciting sustainable ‘sterilizing immunity’ at the mucosal level. Our study uncovers, strong neutralizing mucosal component (NT50 ≤ 50pM), emanating from intramuscular administration of an mRNA vaccine. We show that saliva of BNT162b2 vaccinees contains temporary IgA targeting the Receptor-Binding-Domain (RBD) of SARS-CoV-2 spike protein and demonstrate that these IgAs are key mediators of potent neutralization. RBD-targeting IgAs were found to associate with the Secretory Component, indicating their bona-fide transcytotic origin and their dimeric tetravalent nature. The mechanistic understanding of the exceptionally high neutralizing activity provided by mucosal IgA, acting at the first line of defence, will advance vaccination design and surveillance principles, pointing to novel treatment approaches, and to new routes of vaccine administration and boosting.

**Significance statement:** We unveiled powerful mucosal neutralization upon BNT162b2 vaccination, mediated by temporary polymeric IgA and explored its longitudinal properties. We present a model, whereby the molecular architecture of polymeric mucosal IgA and its spatial properties are responsible for the outstanding SARS-CoV-2 neutralization potential. We established a methodology for quantitative comparison of immunoreactivity and neutralization for IgG and IgAs in serum and saliva in molar equivalents for standardization in diagnostics, surveillance of protection and for vaccine evaluations.

## Introduction

Sterilizing immunity is defined as the ability of the immune system to prevent massive replication and subsequent transmission of a pathogen. Primary infection of some viral pathogens at mucosal surfaces is capable of eliciting sterilizing mucosal and systemic immunity, which is activated in a case of secondary exposure (e.g. enteric Polio- and Rota-viruses; as well as respiratory Influenza virus). Mimicry of certain elements of viral infection by vaccination aims to train the immune system to be tuned for subsequent challenges with the actual pathogen (Holmgren and Czerkinsky, 2005). An ultimate goal of a vaccination campaign besides protection against the disease and death is to achieve a robust sterilizing effect, alleviate the carrier state and interrupt the transmission cycle in the population (Pollard and Bijker, 2021). In this view, vaccine efficiency has several distinct, albeit interconnected aspects – (i) reduction of viral load at the entry site and preventing spread between individuals; (ii) preventing viral spread within the host and expediting virus clearance; (iii) protection from symptoms or reducing disease severity.

During the natural course of viral infections pre-symptomatic and asymptomatic individuals can transmit the virus. In an analogy, a vaccine that protects from the disease does not necessarily achieves the sterilizing effect. Pathogen-targeting IgA at mucosal surfaces is known to correlate with sterilizing immunity, thereby preventing transmission of respiratory and enteric viruses (Blutt et al., 2012; Donlan and Petri, 2020; Lavelle and Ward, 2021; Mostov and Deitcher, 1986; Renegar et al., 2004; Wright et al., 2014; Yu et al., 2021).

Severe Acute Respiratory Syndrome Coronavirus-2 (SARS-CoV-2), the etiological agent of Coronavirus disease 2019 (COVID-19), is a highly contagious and difficult to contain respiratory virus, regardless of disease status and severity, mainly because both asymptomatic and pre-symptomatic individuals are responsible for substantial transmission events (Harrison et al., 2020; Zhou et al., 2020). The two mRNA COVID-19 vaccines, BNT162b2 (Pfizer/BioNtech) and mRNA-1273 (Moderna), has successfully reduced the burden of symptomatic COVID-19, and its more serious outcomes, e.g. (Bruxvoort et al., 2021; Dagan et al., 2021; Haas et al., 2021)). However, the emerging SARS-CoV-2 variants raise concerns as to its long-term protective capability. The immunoglobulin gamma (IgG) responses to natural SARS-CoV-2 infection and the role of IgG against the Spike protein and its Receptor Binding Domain (RBD) in virus neutralization and disease prevention are well established, e.g. (Isho et al., 2020; Pullen et al., 2021; Röltgen et al., 2022; Yu et al., 2021). The IgG response to the vaccine has been thoroughly reported, e.g. (Danese et al., 2021; Levin et al., 2021; Sokal et al., 2021).

IgA is the most abundant immunoglobulin isotype in humans with daily secreted amounts reaching 60 mg/kg/day (Kutteh et al., 1982; Monteiro and Winkel, 2003). IgA plays a key role in the interaction between the immune system and environmental insults to provide mucosal protection, often serving as the first line of defence (Kerr, 1990; Woof and Russell, 2011). Beyond its documented role at mucosal surfaces, IgA is the second most abundant isotype in the blood circulation following IgG, with about 20% of total circulatory immunoglobulin content. Serum IgA is predominantly a monomer, whereas secreted IgA at mucosal surfaces appears in a dimer/polymer form (Kerr, 1990). The IgA dimer, joined through the J chain via disulfide bridges, forms a secretory component (SC-IgA) together with a portion of the polymeric immunoglobulin receptor (pIgR), that is necessary for trans-epithelial secretion (Brandtzaeg, 1981, 2013; Brandtzaeg and Prydz, 1984; Mostov and Deitcher, 1986). Still, in some cases, monomeric IgA can be found at mucosa, and traces of multimeric IgA have been reported in serum as well (Kutteh et al., 1982). Importantly, the mechanistic relationship between mucosal and systemic immunoglobulin responses are not fully resolved (Iversen et al., 2017). Both monomeric and dimeric RBD-targeting IgA elicited by SARS-CoV-2 infection were shown to possess strong neutralizing potential in biological fluids and when tested in a monoclonal Ab setup (Cervia et al., 2021; Sterlin et al., 2021; Wang et al., 2021b; Zeng et al., 2021). However, the immunological characteristics and kinetics of the IgA response, particularly of its mucosal component, upon mRNA vaccination have not yet been deeply investigated, e.g. (Bleier et al., 2021; Danese et al., 2021; Gray et al., 2021; Juncker et al., 2021; Ketas et al., 2021; Matuchansky, 2021; Russell et al., 2020; Wisnewski et al., 2021).

Here we analysed the humoral immune response to BNT162b2 vaccine and detected transitory secretory dimeric IgA, which targets the RBD of SARS-CoV-2 spike in the saliva of vaccinees. We unveiled the powerful neutralizing activity of this humoral component of the mucosal defence and explored its kinetic profile. Furthermore, we established a methodology for quantitative comparison of immunoreactivity and neutralization for humoral IgG and IgA response in serum and saliva in molar equivalents. We submit a universal approach for assessment of antibody response that can be applied globally and will ease standardization in diagnostics and surveillance practices, in decision making in patients’ care, and in comparative vaccine evaluations.

## Results

### Kinetic profiling of circulatory IgG and IgA immunoglobulins in BNT162b2 vaccinees

In the course of monitoring the kinetics of the serological response in a BNT162b2 vaccinated cohort, we noticed that along with a well characterized IgG response toward RBD, a substantial proportion of vaccinees developed a time-dependent accrual of RBD-targeting IgA. To further understand functional aspects of BNT162b2 protection, we studied serum immunoglobulin responses to the vaccine and their kinetic properties. Serum samples were taken from 90 participants (Table S1, cohort details), including pre-COVID-19 cohort, COVID-19 convalescents and vaccinees aged 24 to 75, who received two BNT162b2 doses at a three-week interval (time points included pre-vaccination and follow up till six-month after the first vaccine dose). A more detailed longitudinal follow-up cohort included serum samples (N=76) collected from 18 participants (Table S2, cohort details).

We focused on IgG and IgA directed against RBD region of the viral Spike, as many studies have shown that anti-RBD Abs hamper SARS-CoV-2 entry into host cells by competing with the binding to the host-cell receptor Angiotensin Converting Enzyme 2 (ACE2) (Hoffmann et al., 2020; Ju et al., 2020; Lan et al., 2020; Letko et al., 2020; Li et al., 2005). To measure the antibody response, we produced fully glycosylated recombinant SARS-CoV-2 RBD in a mammalian expression system and used a custom ELISA amenable to quantitative measurements (see Figures S1A-C, methods and supplementary section for details and validation).

Robust anti-RBD IgG and IgA activity was evident in all vaccinees at 10-30 days after the second vaccine dose, *versus* naïve (pre-COVID-19) individuals (Figure 1A). Overall, vaccine-induced anti-RBD IgG was stronger than in convalescents, while the IgA levels were comparable between the two groups, in agreement with recent reports (Cho et al., 2021; Jalkanen et al., 2021; Wang et al., 2021a). This suggests that at least from the quantitative perspective BNT162b2 vaccine prime/boost regimen initiates anti-RBD humoral immune response in circulation comparable to or even stronger than the one observed upon recovery from the natural infection.

**Figure 1:**
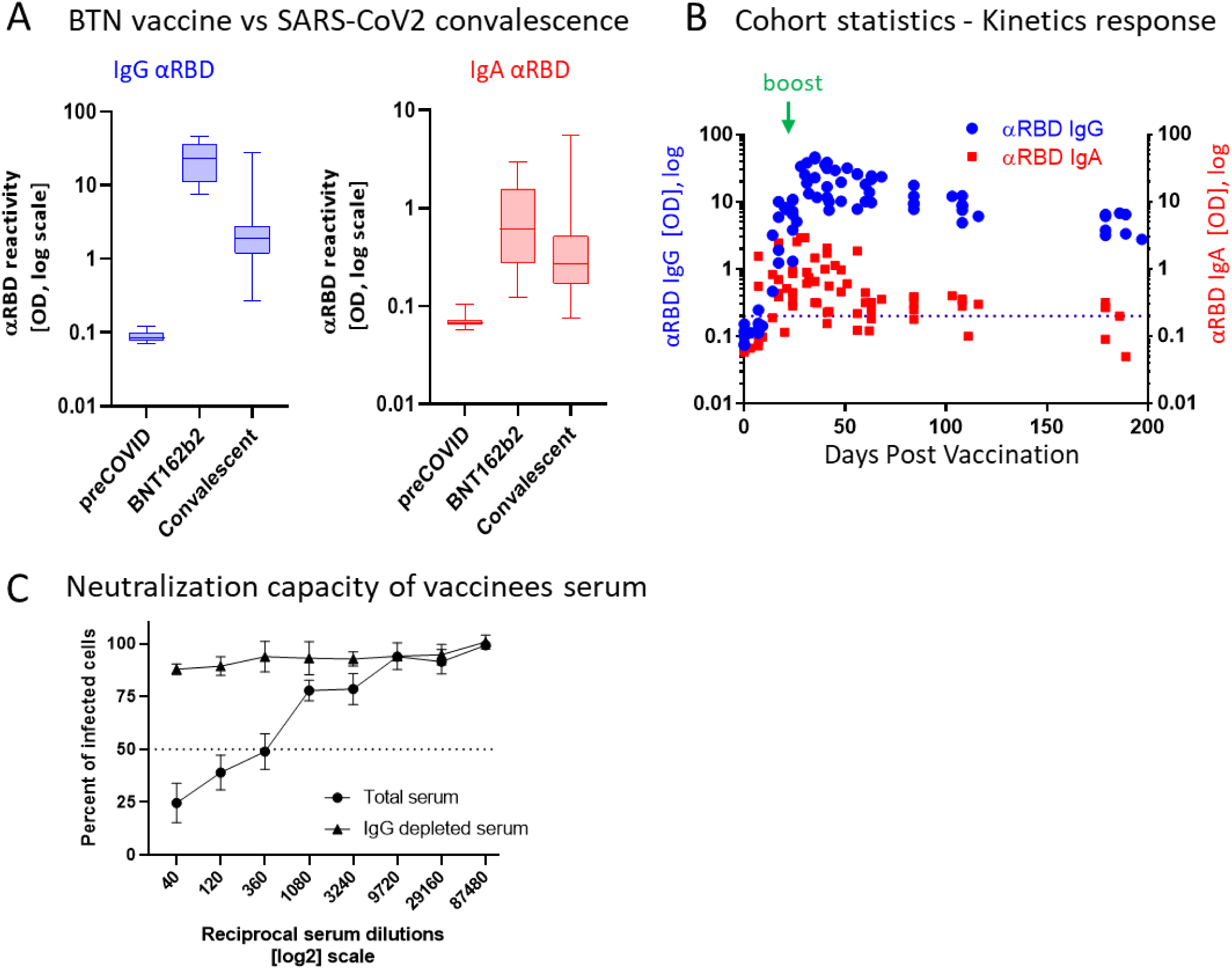
BNT162b2 vaccinees mount serum antiRBD-SARS-CoV-2, IgG and IgA with IgG showing strong neutralisation potential. (A) Independent ELISA measurements of anti-RBD IgG and of anti-RBD IgA in serum samples collected from pre-COVID (N=51), BNT162b2 vaccinees (N=17) and post-COVID-19 (N=22) convalescents, as indicated. Convalescent samples were collected within two months post-recovery. BNT162b2 vaccinees samples represent the peaks of individual responses. (B) Quantitative kinetic profile of anti-RBD IgG (blue) and IgA (red) in serum sampled (N=76) in the vaccinees cohort (N=18), plotted as a function of days, post first vaccine dose. See Table S1 for cohort and sampling details. Independent ordinate axes for IgG (left, blue) and IgA (right, red) highlight the restricted, relative nature of the comparison between isotypes in this experiment, as discussed in the text, see also Figure 2 for subsequent developments. Green arrows indicate timing of the second vaccine dose (the boost). (C) Serum neutralisation assessed by SARS-CoV-2-Spike pseudotyped VSV-GFP-ΔG reporter assay on Vero-E6 cells. Neutralisation is expressed as a percentage of pseudovirus-infected green cells without serum (total infection = 100%). Percentage of neutralisation by sera of pool of four individual vaccinees (see the corresponding anti-RBD IgG and IgA values and times post vaccination in Figure S1) are plotted as a function of the reciprocal values of sera dilutions displayed on a log2 scale, as indicated (filled circles, total serum, NT50 is reached on average at the dilution of ∼1:360, extrapolated by cross-section with the dashed line. Contribution of IgG to serum neutralisation is evaluated by the depletion of the IgG isotype using anti-IgG specific magnetic beads (triangles). Results of three experimental repeats are represented.

Next, we carried out a detailed time course analysis of the serological response among the vaccinees. Notably, the fact that the majority of commercially available SARS-CoV-2 antibody assays, as well as our results presented in Figure 1B, use arbitrary unit values, impedes the capability to directly compare anti-RBD IgG and IgA levels in terms of molecular stoichiometry. Therefore, we used two different ordinate axes to represent IgG and IgA arbitrary values and only relatively superpose the respective shape and durability of the two isotypes. The magnitude of the serological IgA response among vaccinees was significantly more scattered and overall showed less steep increase than that of the IgG (Figure 1B), suggesting a higher variability of the vaccine in induction of IgA isotype in the circulation. Monitoring of the circulatory levels for six months post-vaccination in a cohort subset revealed a decline of anti-RBD IgG and IgA (Figure 1B, see also Figure S1D for violin plot representation of categorized periods).

In order to assess the specific functional contribution of IgA and IgG in serum we performed neutralization analyses using a reporter assay in Vero E6 cells, based on SARS-CoV-2 spike-pseudotyped Vesicular Stomatitis virus (VSV). First, we selected a pool of sera from four vaccinated individuals with significant anti-RBD IgA and IgG levels and depleted total IgG molecules. Figure S1E shows the complete drop in total IgG levels upon depletion, as measured by sandwich ELISA (depicted in Figure 2B). While the original sera pool showed the half neutralization capacity (NT50) at the dilution of ∼1:360, the IgG depleted pool resulted in a complete loss of neutralization (Figure 1C). This indicates that the vaccine-elicited IgG is the functionally predominant neutralizing isotype in blood circulation in accordance with previous findings (Cho et al., 2021; Turner et al., 2021; Wang et al., 2021c).

**Figure 2:**
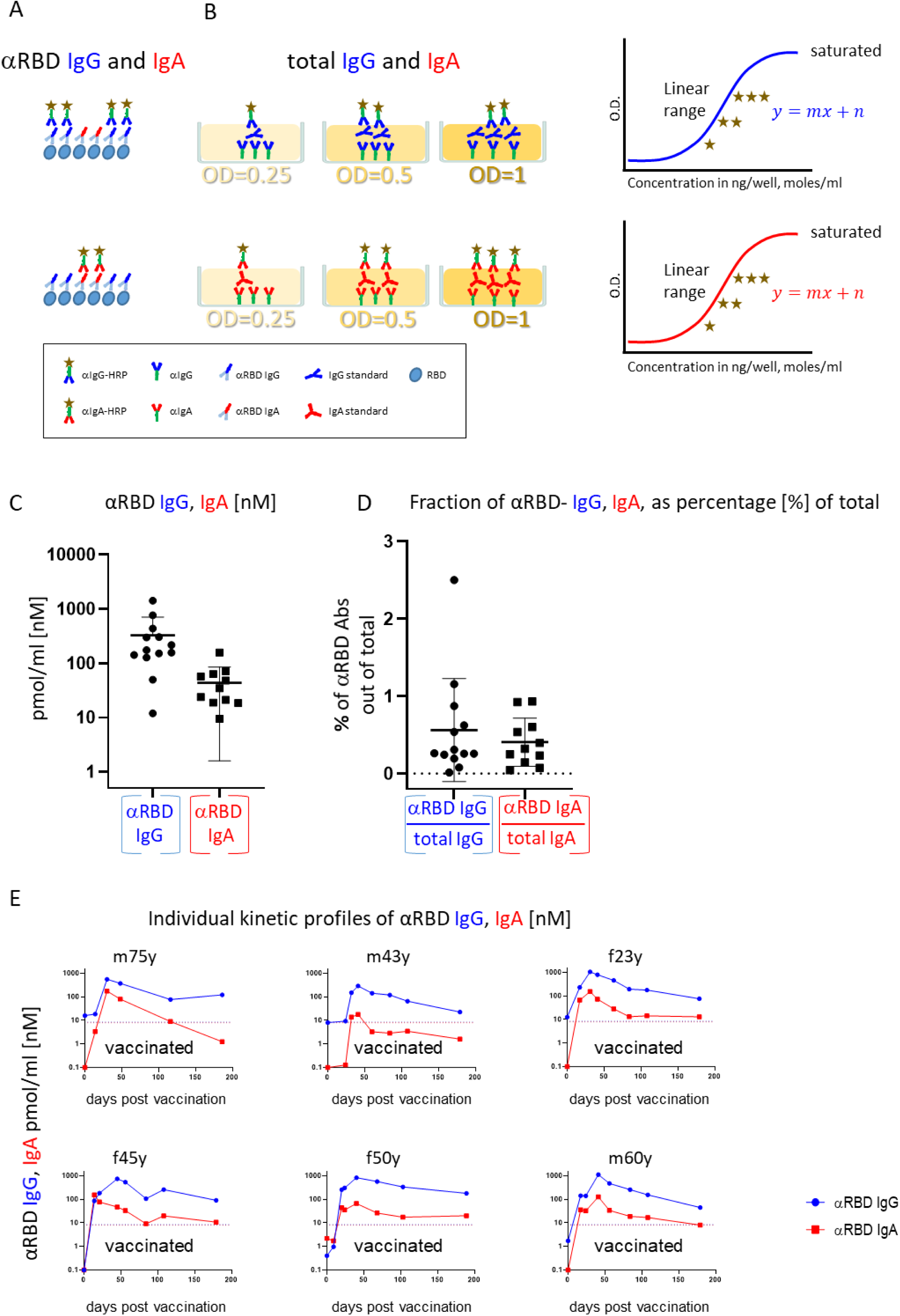
Quantitative ELISA measurement of anti-RBD IgG and IgA content in biological fluids. (A) Schematic representation of detection of anti-RBD IgG or IgA by indirect ELISA using isotype-specific HRP-conjugated secondary antibodies. OD values are not directly comparable between the isotypes because of the use of different secondary antibodies. (B) Schematic representation of sandwich capturing ELISA for selective quantification of total immunoglobulin isotypes (IgG vs IgA). We introduce pure IgA and IgG commercial references to transform the OD values to their molar equivalents, using standard dilution curve in capture ELISA format. Implementing such a standard in every experiment allows determining the antigen specific and total molar amounts of each isotype within the linearity range. We assume average molecular weight (MW) of IgG= and IgA=160kDa in circulation. (C) Molar measurement of anti-RBD IgG and IgA implementing the methodology described in panels A and B depicts stoichiometric ratios between the antigen-specific isotypes. (D) Percentage of antigen-specific anti-RBD out of total immunoglobulin isotype, as indicated. (E) Individual longitidual profiles of anti-RBD IgG (blue) and IgA (red) monitored in six vaccinated individuals up-to 7 month post vaccination are inferred as picomole per ml of serum. Gender and age of vaccinees are indicated on each plot.

### Measuring absolute and proportional amounts of vaccine-elicited IgA and IgG

BNT162b2 vaccine-elicited circulatory anti-RBD IgA has drawn our attention, particularly due to the crucial importance of IgA in providing the ultimately desired mucosal defence.

As experience with intramuscular RNA vaccination is limited, especially with respect to IgA response overall, and in particular at the mucosal interface, we decided to explore the role of secretory IgA. One notable obstacle in the functional assessment of the role of IgA in both circulation and mucosal surfaces is the inability to quantify and compare circulatory and mucosal IgG *versus* IgA. In our view, this is of utmost importance for SARS-CoV-2 studies, in particular due to the current need for a universal absolute measure of humoral response at different physiological sites (WHO/BS/2020.2403). We tackled this obstacle at three levels: (1) Serial dilutions of the serologically evaluated biological fluids to empirically determine the linear confidence range in immunoassays. (2) Implementation of pure human IgG and IgA fractions to create a calibration curve to convert Optical Density (OD) of secondary detection into absolute quantitative units (e.g. moles) independent of the secondary Ab conjugates, e.g. (Kannenberg et al., 2022; Wang et al., 2021a). (3) Evaluation of the specific contribution of anti-RBD IgA or IgG by determining their proportion out of total immunoglobulins of the same isotype in a given biological fluid and subsequent functional assessment of the isotype-specific depletion. We used serum samples of vaccinated individuals, described in Figure 1B to establish and validate such measurements (see table S1 for the sub-cohort details).

In a typical ELISA used to measure RBD antibodies, we coat the plate with RBD and subsequently react it with the relevant biological fluids. RBD-reacting Abs of all isotypes are captured, while IgAs and IgGs are then differentially revealed by the corresponding isotype-specific secondary Abs (Figure 2A). Using commercial pure human IgG and IgA standards with defined concentrations, we established ‘OD-to-mole’ transformation (Figure 2B). To this end we use an ELISA setup, measuring the total IgG and IgA populations rather than antigen-specific subsets. In this case, the plate is first coated by capturing isotype-specific Abs and then the defined amounts of the reference isotypes are entrapped and revealed by the isotype-specific Ab-HRP (Horseradish Peroxidase) secondary conjugates. By introducing such a standard in our experimental routine, we could quantitatively relate OD to the absolute amount of captured immunoglobulins (Figure S2A-C). Figure S2D shows the specificity of isotype capturing and detection, with no apparent cross-reaction.

We next applied this approach to evaluate molar concentrations of RBD-targeting IgG and IgA in the serum of vaccinees. Figure 2C quantitatively shows that the majority of individuals produced 200-1000 pmol/ml (nM) of RBD-targeting IgG *versus* 30-200 pmol/ml (nM) of RBD-targeting IgA. Our approach allowed the determination of the proportion of RBD-specific Abs out of the total amount of immunoglobulins of the given isotype in serum, (normalised, proportional formula 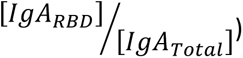 (Figure 2D). While absolute quantities of RBD-specific IgG strongly dominate over the corresponding IgA in serum, the normalised fractional quantities were comparable and comprise 0.5 % of IgG and 0.4 % of IgA in circulation (Figure 2D, graphical representation and Table S4, sub-cohort data). This near equal proportional representation of both isotypes toward the RBD antigen upon intramuscular mRNA vaccination, suggests a similar frequency of class-switch events. In the next series of experiments, we monitored IgA and IgG kinetics expressed in pmol/ml values of anti-RBD in the serum of six individual vaccinees, allowing stoichiometric comparison (Figure 2E). All the individuals exhibited predominant IgG response that peaked around 40d post vaccination and gradually declined during six months. In contrast, IgA responses were more variable (Figure 2E). In all our measurements, the circulatory IgA picomole values were lower and with shorter duration than those of IgG, similar to other vaccine instances and upon natural immune responses to infections (Salonen et al., 1985). Of note, we considered the IgA-monomer in serum for our molar transformations, as circulatory IgA is most commonly monomeric lacking the secretory component, while dimeric IgA is mostly found at mucosal surfaces and in mucosal secretions (see also Figure 4).

### Robust anti-RBD IgA response in saliva of vaccinees

Given the substantial IgA amounts in serum elicited by the vaccine, and the well-established role of secreted IgA in providing mucosal immunity, we asked whether RBD-reactive IgA can be detected in resting saliva of vaccinees. Saliva samples (N=82) were obtained from 33 participants, aged 20-75 (see Table S5, cohort details).

First, we confirmed that the vast majority of total immunoglobulins in saliva detected by our quantitative ELISA were of the IgA isotype, in agreement with the well-characterized humoral repertoire of the salivary milieu (Figure S3). Next, we turned to quantify the RBD-specific IgA in saliva samples collected at different time points post-vaccination, as indicated (Figure 3A). Anti-RBD salivary IgA response was rather variable between individuals, akin to its variability in serum. There was a time-dependent increase in anti-RBD IgA, peaking at about 3-4 months post-vaccination and then decreasing steeply at about 5 months post-vaccination, dropping to the levels of naïve individuals by the end of 6 months. Notably, the duration of mucosal anti-RBD IgA was significantly extended when compared to circulatory IgA, suggesting involvement of certain aspect of mucosal immunological memory (see discussion).

**Figure 3:**
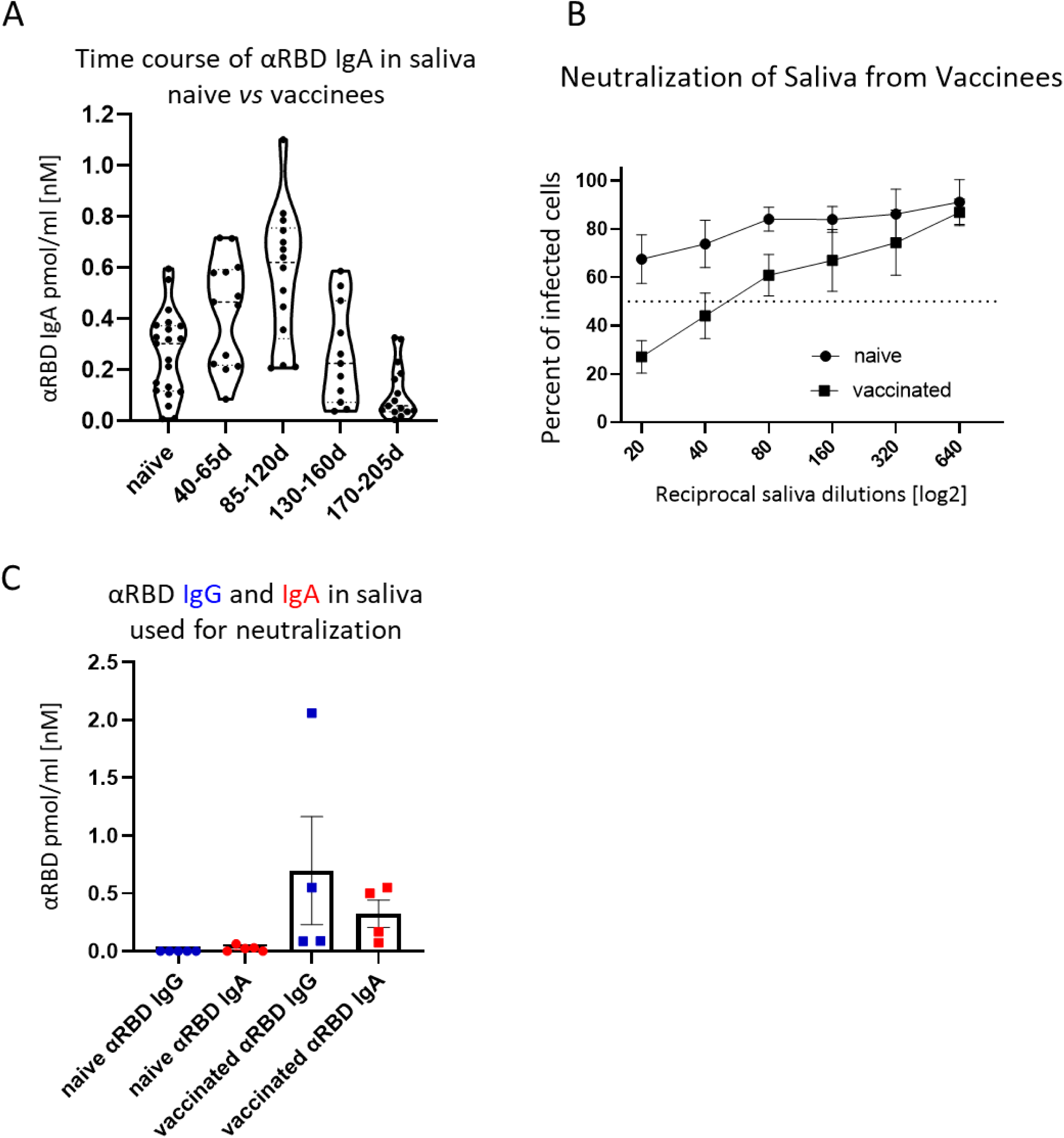
Detection of anti-RBD IgA in resting saliva of BNT162b2 vaccinees and characterization of its neutralizing potential. (A) Longitudinal assessment of molar quantities of anti-RBD IgA in saliva of vaccinees compared to naïve individuals is presented in picomole per ml and time-categorized as indicated. The molar expression in saliva is corrected to bi-valence for the convenience in comparison to circulatory immunoglobulins (B) Saliva neutralisation assessed by SARS-CoV-2-Spike pseudotyped VSV-GFP-ΔG reporter assay on Vero-E6 cells. Neutralisation is expressed as a percentage of pseudovirus-infected green cells without incubation with saliva (total infection = 100%). Percentage of independently measured neutralisation by five naïve individuals versus five vaccinees are plotted as a function of the reciprocal values of sera dilutions displayed on a log2 scale, as indicated. Each neutralisation curve was tested in three biological replicates. Standard deviation represents difference between individuals in each group. The NT50 of vaccinees saliva is achieved on average at the dilution of 1:60, extrapolated by cross-section with the dashed line. The specific neutralisation NT50 value is reached at dilution of 1:20 and represents ‘vaccine-added’ neutralisation, corrected to the basal innate neutralisation of naïve individuals, that is probably the consequence of innate proteolytic and mucus (lectin) presence in naïve saliva. (C) The values of anti-RBD IgA and IgG in picomole per ml of five saliva samples, used in neutralisation assay described in panel B are shown.

The presence of anti-RBD immunoglobulins in saliva is rather encouraging, though the question remains as to its ability to prevent virus entry. Using the VSV-GFP-SARS-CoV-2-Spike pseudotype neutralization assay, a strong concentration-dependent neutralizing activity of saliva from vaccinees was discovered (∼NT50 1:60) (Figure 3B, squares, five vaccinees samples). This value is significantly higher than the basal background neutralizing activity of saliva from naïve individuals (Figure 3B, circles). The background neutralizing activity of saliva from naïve individuals may stem from basal innate antiviral properties of naïve saliva (e.g. proteolytic digestion and lectin properties). For the sake of sterility in neutralization assay, we used pre-diluted saliva samples that were cleared by centrifugation (12,000g, 5min) and subsequent size filtration (0.22µm) (see Figure 3C for the ELISA of clarified saliva samples used in Figure 3B). The solubility of IgA molecules in saliva is often a matter of concern due to the viscous-colloid, mucus state. We confirmed quantitative recovery of solubilized saliva IgA by comparing pre- and post- centrifugation and filtration samples by quantitative ELISA (Figure S3B and Tables S6).

### Depletion of IgA from saliva samples of vaccinees completely abrogates neutralization activity

Given the IgA prevalence in saliva, we turned to evaluate its functional contribution to neutralization (Figure 4A). We generated a pool of five saliva samples from vaccinees, and subsequently depleted either the IgA or IgG molecules (Figure S4). Depletion of IgA, but not IgG, resulted in the loss of neutralization (Figure 4A). This result confirms that the strong neutralizing activity of vaccinees is attributed to IgA.

**Figure 4:**
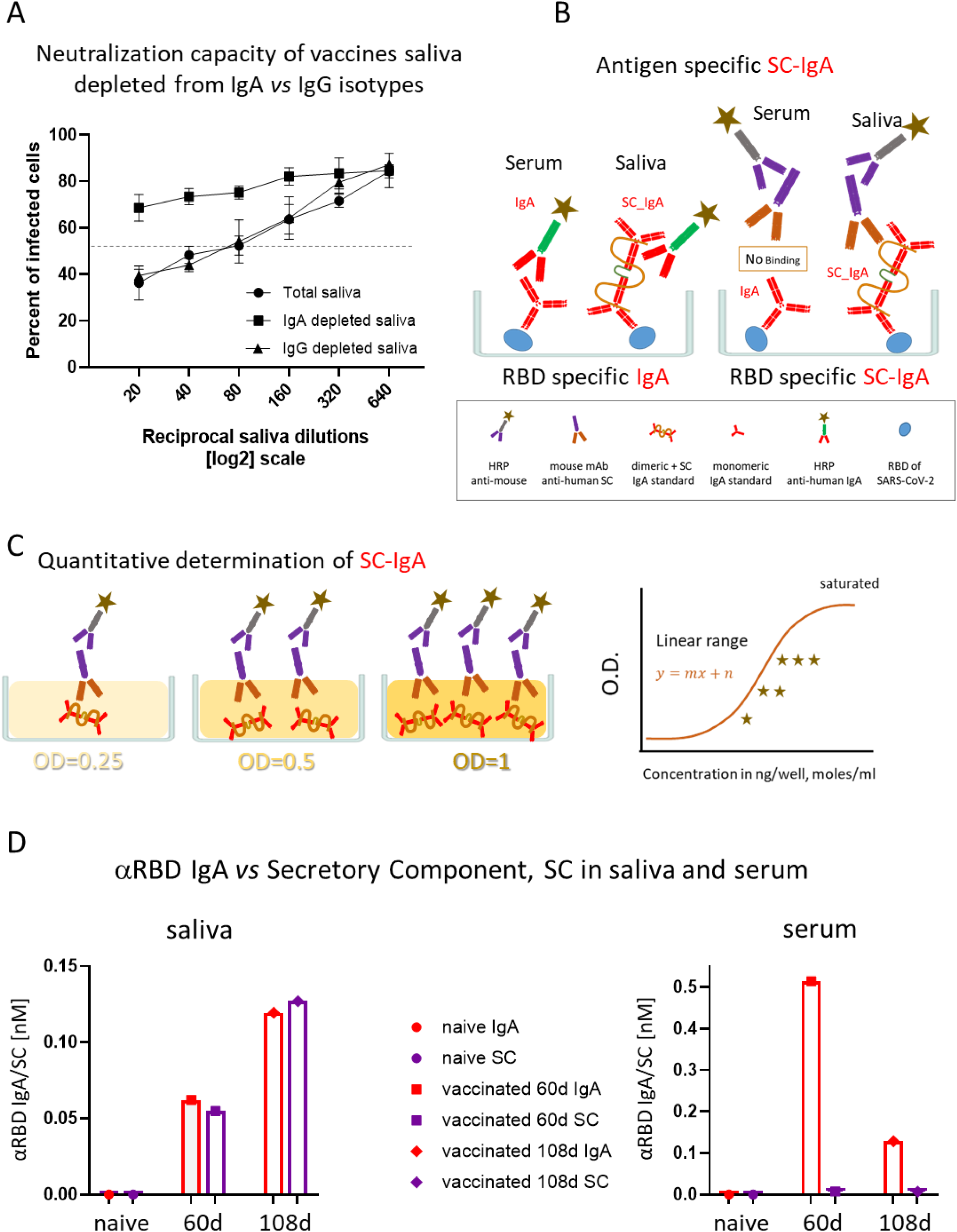
The association of salivary anti-RBD IgA with the secretory component governs the prominent neutralisation activity in vaccinees. (A) Depletion of IgA from saliva samples of vaccinees completely abrogates the specific neutralisation activity of vaccinees saliva. Saliva neutralisation was assessed by SARS-CoV-2-Spike pseudotyped VSV-GFP-ΔG reporter assay on Vero-E6 cells. The magnitude of neutralisation is expressed as a percentage of pseudovirus-infected green cells without incubation with saliva (total infection = 100%). Percentage of measured neutralisation by saliva pool of five vaccinees is plotted as a function of the reciprocal values of the saliva dilutions displayed on a log2 scale. The corresponding anti-RBD IgA and IgG values of individual saliva samples chosen for the pool show clear quantitative dominance of IgA, Figure S4. The NT50 of saliva pool is reached at the dilution of 1:60 (extrapolated by the cross-section with the dashed line). Depletion of IgA results in abrogation of vaccine-induced neutralisation activity (squares), while IgG depleted saliva pool perfectly coincides with the non-depleted pool (triangles *versus* circles). Depletion is achieved using anti-IgA and anti-IgG specific magnetic beads. Analyses of completeness of isotype depletion and of its specificity are presented in Figure S4. Results of three experimental repeats are represented. (B) Schematic outline of the detection of anti-RBD IgA in serum and SC-associated anti-RBD IgA in saliva samples. Illustrated are the expected differences between the circulatory monomeric IgA and the salivary mucosal dimeric/polymeric IgA, covalently bridged by J-chain and associated with pIgR. Left panel shows non-discriminative detection of both isoforms by anti-IgA secondary HRP-conjugate. Right panel shows the selective quantitative determination of dimeric secretory IgA in saliva, but not in the serum, using anti-SC mouse monoclonal Ab, followed by anti-mouse secondary detection. (C) Molar quantification of dimeric secretory IgA. We introduce reference standard using commercial secretory dimeric IgA purified from human colostrum to transform the OD values to their molar equivalents. We assume average molecular weight (MW) of dimeric secretory IgA = 424 g/mole. (D) Analysis of dimeric anti-RBD SC-IgA in saliva (upper panel) *versus* monomeric anti-RBD IgA in serum (lower panel) post vaccination, measured by quantitative ELISA. The molar expression in saliva is corrected to bi-valence to simplify the comparison to circulatory immunoglobulins.

Salivary IgA, similar to all mucosal IgA forms, is produced in a dimeric, tetravalent form, as opposed to bivalent IgA and IgG monomers found in the circulation. To verify whether this is the case in the saliva samples of vaccinees, we employed anti-Secretory Component (SC) quantitative ELISA, measuring molar values of total and anti-RBD secretory dimeric IgA (experimental flow is depicted in Figures 4B and C, see details in Supplementary). We compared a pool of four saliva samples from naïve individuals to the two pools collected from vaccinees at two and three months’ post vaccination. In parallel, we analysed the corresponding serological samples. Figure 4D demonstrates that anti-SC reveals RBD-targeting reactivity solely in the saliva of vaccinees and not in their sera. At 60d post vaccination overall anti-RBD IgA reactivity dominated in serum, while the SC-associated anti-RBD form predominated in saliva, in accordance with the strikingly eminent specific neutralization potency of salivary IgA (Figures 3C and 4C). In contrast to a sharp decline of anti-RBD IgA in serum at day 108 (3.5 months) post vaccination, the salivary IgA presence and its association with the SC were mounting, in line with data shown in Figures 1B and 3A. We conclude that anti-RBD IgA in saliva of vaccinees originates from *bona fide* transcytotic secretory pathway, validating its dimeric nature (Figure 4D).

### Strong neutralizing activity of salivary polymeric SC-IgA *vs* monomeric IgG and IgA in serum

To establish the specific neutralization potency of the anti-RBD immunoglobulins, we normalised their relative NT50 values to their actual [nM] concentration in the respective fluids (Figure 5-I). This pointed to a two orders of magnitude advantage of saliva anti-RBD IgA (NT50 = 0.02-0.05nM), *vs* serum anti-RBD IgG (NT50 ∼1nM). We hypothesize that the remarkable neutralization potency of polymeric salivary IgA (relative to monomeric serum IgG) may stem from a combination of (i) the increased avidity of multivalent binding and (ii) a geometrical fit between dimeric IgA and the SARS-CoV-2 spike protein trimer as presented on the surface of virions.

**Figure 5:**
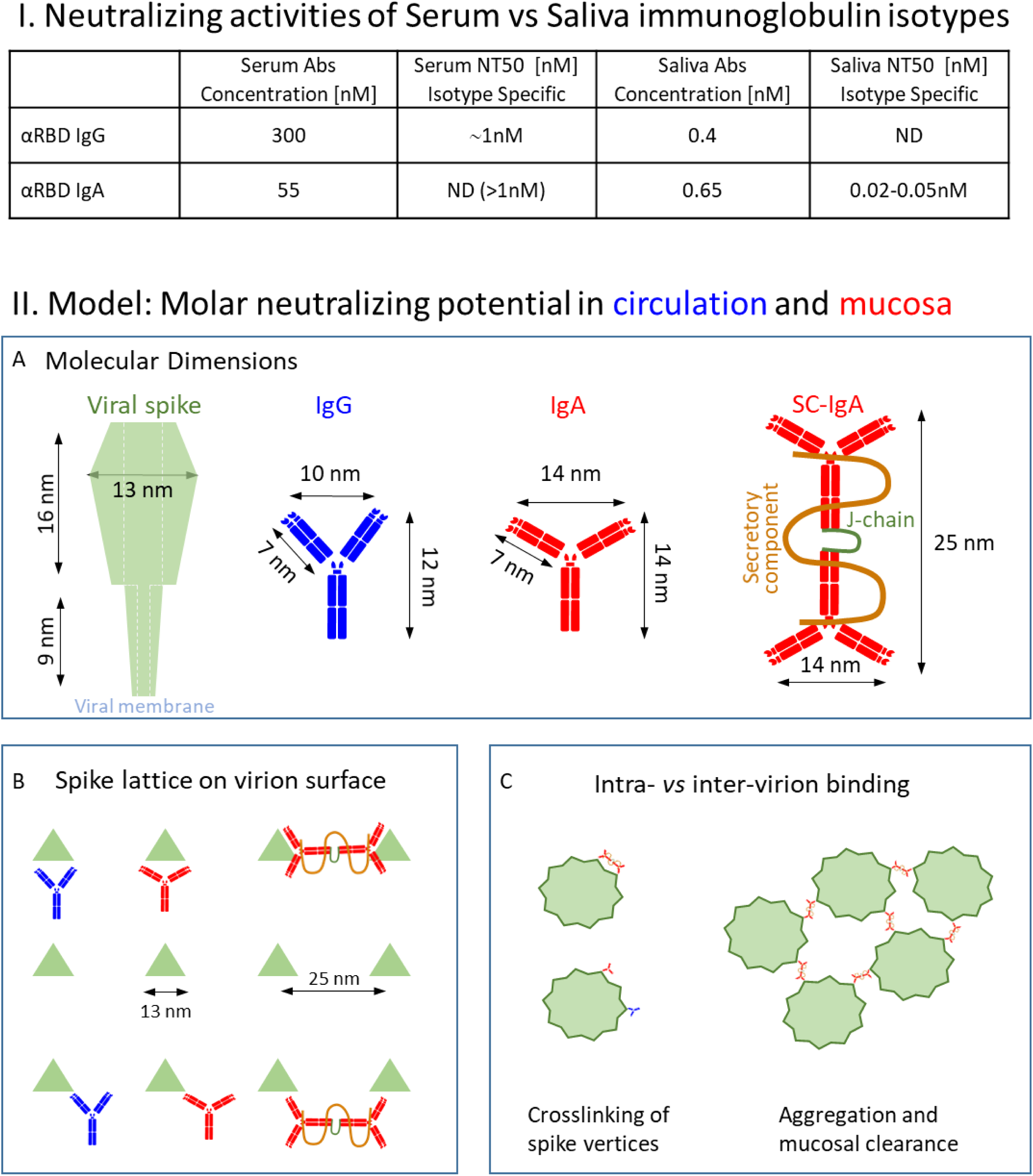
(I) Strong neutralizing activities of saliva anti-RBD SC-IgA vs serum immunoglobulin isotypes. Quantitative, molar measurements of anti-RBD immunoglobulin content in saliva and serum allow the evaluation of the specific neutralizing activity expressed as NT50 per [nM] of anti-RBD IgA and IgG per in saliva and serum, respectively. To calculate these values we normalised NT50 expressed in dilutions of serum and saliva (Figures 1 and 3) to the molar concentration in saliva and serum, respectively [nM]. NT50 value for salivary IgA was calculated based on average NT50 dilution of 1:20, upon normalization to basal inhibitory activity of naïve saliva (see Figure 3B). NT50 value for salivary IgA was calculated based on average NT50 dilution of 1:20 anti-RBD IgG, average dilution of 1:300), (Figure 5-I). The plausible mechanisms behind such stark, two orders of magnitude difference in NT50 between salivary and serum immunoglobulins are addressed in the ‘Thought experiment’ described below and summarized in the model presented in Figure 5-II. **(II) Model: Molar neutralizing potential in circulation and mucosa - ‘*GedankenExperiment*’ to explain protective outcome.** (A) Known dimensions of SARS-CoV-2 spike protein trimer next to dimensions of circulatory and mucosal immunoglobulins. The extension and flexibilities of IgA arms are illustrated by wider angularity of the Fabs for the monomer. The longitudinal extension of the dimeric SC-IgA is represented in the right panel. (C) Lattice density of trimeric spike vertices, represented by green triangular surface projections of a viral antigen and their coverage by immunoglobulins are shown (see text for details). (D) Illustration of protective outcomes: SC-IgA versus IgG surface interaction areas. (E) Illustration of the interaction modalities in the context of the virus particle: intra-virion binding of SC-IgA and monomeric IgG and IgA (left panel) and inter-virion aggregation by SC-IgA (right panel).

Whereas the avidity components of multivalent binding and neutralization are well studied (Laursen et al., 2018) in viral infections, the subject of complementarity between a virion lattice and the immunoglobulin isotype is less explored. The simplified view of the molecular dimensions of SARS-CoV2 spike and of the studied immunoglobulin isotypes are presented in Figure 5-IIA (molecules are drawn schematically with respect to their proportional scale). Since the surface glycoprotein lattice is sparse (e.g. majority of trimeric spike vertices are 20-25nm apart) (Klein et al., 2020; Yao et al., 2020), circulatory IgGs and IgAs might bind to only single glycoprotein spike restricted by their Fab arm spread of (10-14nm). In contrast, dimeric SC-IgA can concomitantly capture two glycoprotein spikes due to its 25nm-longitudinal extension, thereby more efficiently covering – ‘mantling’ the virion surface. In this view, the ‘mantling efficiencies’ of dimeric secretory IgA are reminiscent of a mythical warrior (*Sanskrit* ‘*Virabhadra* - **}{**‘) and might by far outperform the restricted capabilities of IgG (Figure 5-IIB and C, left).

Figure 5-IIC illustrates the whole-virion perspective and outlines an additional layer of plausible protection, whereby SC-IgA induced inter-particle oligomerization is promoted by the extended and elastic tetravalent branches. Such a situation, fortified by a plethora of polyclonal species, may appear as highly prominent *in vivo*. In oral immunity, the aggregation of exogenous particles is known to promote mechanical clearance of invaders from mucosal surfaces (Bustamante-Marin and Ostrowski, 2017; Nail et al., 1969). The consequences might become even more significant in the case of a higher degree of multimerization that was recently observed for human intranasal IgAs. Intranasal vaccination with *Influenza* virus in humans has revealed extreme potency and ‘neutralization-breadth’ toward variant strains. This powerful protection was attributed to nasopharyngeal SC-IgA with the elevating superiority of the multimeric states: dimers, trimers, tetramers and even higher order oligomers (Suzuki et al., 2015).

Our findings, in conjunction with the ‘*GedankenExperiment*’ (Figure 5-IIB and C) of interaction modalities between surface SARS-CoV-2 lattice and mucosal dimeric IgA *vs* monomeric IgG and IgA in blood circulation, highlight the importance of implementing lattice design to improve the spatial surface-mimicry in the next-generation subunit vaccines. In this respect, the mRNA based vaccine may have had an unexpected benefit by enabling the host-cell to present the natural arrangement of SARS-CoV-2 spike membranal lattice upon its expression.

Overall, our results demonstrate that the BNT162b2 vaccine induces a five-month transient accrual of salivary anti-RBD IgA, extending beyond the time frame of detectable circulatory IgA, putting forward a basis for the establishment of mucosal memory. We suggest that the polymeric origin of the salivary IgA molecules forms the basis for the remarkably high specific neutralizing activity found in the BNT162b2 vaccinees’ saliva, compared to serum IgG. Salivary anti-RBD IgA may represent a more general nasopharyngeal humoral component of mucosal protection.

## Discussion

Our study reveals a previously undescribed mucosal component resulting from the intramuscular administration of an mRNA vaccine. We unveil that saliva of vaccinees contains transitory anti-RBD dimeric secretory IgA (Figures 3A, 4D) with strong neutralizing activity (Figure 3B, Figure 5-I), possibly explained by its tetravalent nature. We show that this polyvalent IgA is the main mediator of potent neutralization activity in the vaccinees’ saliva, remaining unchanged following IgG depletion. Accordingly, vaccine-induced neutralization was completely abolished by depleting salivary IgA. In contrast, IgG was the predominant neutralizing isotype in serum since its removal resulted in loss of neutralization. Intriguingly, and contrary to the situation in saliva, residual serum IgA was devoid of measurable neutralization activity, despite its significantly higher concentration - about 30-fold higher content in serum *vs* saliva (when valence differences are accounted, see Methods). The unique feature of mucosal IgA is its association with the joining J-chain that bridges two iso-clonal immunoglobulin molecules upon their synthesis in secreting B-cell and with the SC that mediates trans-epithelial delivery and extends their lifespan in the highly hydrolytic mucosal environment. The functional and mechanistic impacts of such association in terms of avidity and stereochemical properties are discussed below. Epitopic repertoire of salivary and serum IgA may also differ due to affinity maturation driven somatic hypermutations of Nasopharynx Associated Lymphoid Tissue (NALT) resident B-cell clones. Nevertheless, our NT50 molar measurements, intrinsically normalised to the binding reactivity values, assessed in ELISA, strongly argue in the favour of superior neutralization by salivary IgA due to its polymeric origin.

Anti-RBD IgA remained present in saliva for an extended period of time after vaccination (it peaked at 2-4 months and vanished only 5-6 months post-vaccination), significantly outliving serum anti-RBD IgA (Figure 4D). While analysis of the sustained immunological memory mediated by NALT and Broncho-Alveolar Associated Lymphoid Tissues (BALT) is pending, one might wonder about the possible impacts of systemic or even locally applied mucosal boosts (Lapuente et al., 2021; Tiboni et al., 2021). Since the presence of mucosal IgA and their functional importance in the recovered individuals are now well characterized(Cervia et al., 2021; Sterlin et al., 2021), it might be interesting to monitor the mucosal-effect of post-recovery vaccine boost. The dynamic epidemiological reality, however, is often more complex, given the antigenic diversity of the rapidly emerging SARS-CoV-2 variants. In this view, adaptation of mucosal boosts to emerging variants may be considered in the future.

It is yet not clear whether BNT162b2 mRNA vaccination provides temporary sterilizing immunity, in addition to its proven capacity to ward off severe disease. ‘Sterilizing immunity’ - is crucial for the capacity to break the spread of SARS-CoV-2 and reduce the emergence of new variants. The recent delta-variant wave in Israel where the majority of population has been vaccinated by BNT162b2 approximately 5-6 months prior to the wave spread(Goldberg et al., 2021), coincidentally correlates with our finding of drop in salivary IgA post-vaccination (Figure 3A). Whether waning immunity at the population level had a causative relationship with the drop of systemic anti-RBD IgG or mucosal anti-RBD IgA, and their relative involvements, remain to be investigated. Mucosal IgA are indeed often transient even when induced by natural mucosal invaders, however the immunological memory at the *lamina propria* may reside in place ensuring an inducible defence. It remains yet unknown whether such mucosal humoral memory endures in the vaccinees or in recovered patients and if so, to which extent it reacts to avert spread of the infection. In this vein, recent studies in mice demonstrate feasibility of adenovirus vectored intranasal boosts to achieve complete SARS-CoV-2 protection by (1) inducing high level mucosal neutralizing IgA and (2) stimulating NALT resident memory T cells, when administered after primary mRNA or plasmid DNA intramuscular vaccination (Lapuente et al., 2021).

The ideal vaccine is aimed to provide a perfect mimicry of the natural infection route and as such, would train the immune response to situate its guards ‘*en place*’. The well-studied poliovirus case with the known difference between Inactivated Polio Vaccine (IPV) and Oral Polio Vaccine (OPV) exemplifies the importance of such mimicry for providing the sterilizing immunity (Hird and Grassly, 2012; Onorato et al., 1991). Many more recent studies also demonstrate that the mucosal route of vaccination provides such beneficial protection against respiratory and digestive-tract virus and bacterial infections, including influenza and rotavirus and even SARS-CoV-2 (Hamajima et al., 2002; Hellfritzsch and Scherließ, 2019; JONES et al., 2006; Kurono, 2021; Lapuente et al., 2021; Lavelle and Ward, 2021; Perrone et al., 2009; Terauchi et al., 2018; Yang and Varga, 2014). However, formulating immunogenic, broad, and safe ‘subunit’ or ‘inactivated’ mucosal viral vaccine, capable of eliciting long-term efficient and balanced mucosal-*plus*-systemic protections, remains challenging. Several non-mucosal vaccines were shown to induce the mucosal component of protection (Su et al., 2016), whereas intramuscular DNA vaccines widely studied in animal models are known to possess such capability (Kathuria et al., 2012; Taylor et al., 2005). Intramuscular lipid mRNA formulation in human setting and its mucosal aspects including functional protection, have not yet been widely characterized. Though not tested here, we assume that the elicited mucosal humoral immunity might not be restricted to saliva, and afford broader mucosal protection, extending to: (1) the nasopharyngeal niche elicited by NALT, (2) the lower respiratory Broncho-alveolar mucosa brought about by BALT and even (3) the gastrointestinal tract mediated by GALT. In accordance with this assumption, recent reports indicate presence of anti-SARS-CoV-2 IgA in breast milk of BNT162b2 vaccinated women (Low et al., 2021; Rosenberg-Friedman et al., 2021). Additional support, comes from the recent study by Chan *et al*, who compared two different SARS-CoV-2 parenteral vaccine platforms approved for emergency use in Hong Kong for their ability to induce neutralizing IgG/IgA in serum versus nasal epithelial lining fluid (NELF): CoronaVac (inactivated virus vaccine) and Comirnaty (mRNA vaccine) (Chan et al., 2021). Intriguingly, Comirnaty induced anti-spike neutralizing IgA response detected in nasal epithelial lining fluid, while a similar response was not observed in CoronaVac vaccinees, highlighting the mucosal capabilities of mRNA based *vs* inactivated vaccine (Chan et al., 2021).

In spite of the importance of IgA for protection against pathogens, a certain fraction of the human population is characterized by IgA deficiency (Brandtzaeg et al., 1999; Morawska et al., 2021). Only 10-15% of IgA deficient individuals are susceptible to recurrent sino-pulmonary and gastrointestinal infections/disorders, while the vast majority remains asymptomatic and are often incidentally identified among healthy blood donors. In many cases, IgM appears to compensate the deficiency by replacing IgA at mucosa, as it reacts with pIgR and can be transcytosed to mucosal surfaces (Brandtzaeg et al., 1999). Whether mRNA vaccines boost mucosal IgM as well in such instances of IgA deficiencies remains to be explored.

The mechanism of eliciting the mucosal humoral component by an mRNA vaccine remains mysterious, but the presence of the SC points to transcytotic origin of the polymeric isoform in saliva, generated by the dedicated B-cells, situated at the *lamina propria* (Pilette et al., 2001). Local antigenic stimulation may originate from either the lymphatic drain of the spike protein produced at the site of injection, or from the trafficking to the *lamina propria* of the mRNA itself, being subsequently expressed by either NALT, GALT or the epithelial cells. Analyzing local MHC-I *versus* MHC-II T-cell responses could help in distinguishing between the two scenarios (Heinz and Stiasny, 2021; Rijkers et al., 2021), though potential cross-presentation by dendritic cells may complicate the analysis (Joffre et al., 2012). Such hypothetical delivery of mRNA or of the expressed spike-antigen to mucosa and its potential immunological impact are intriguing and worth a detailed investigation. Plausible routes could involve: (i) a lymphatic drain facilitated by liposome-directed targeting or (ii) a natural exosome-mediated delivery route to distal anatomical sites, or (iii) migration of antigen presenting cells from the site of expression to secondary lymphoid organs. Mechanisms behind such putative scenarios have been previously suggested (Bogunovic et al., 2009; Qiu et al., 2018; Raposo et al., 1996; Théry et al., 1999; Valadi et al., 2007; Zitvogel et al., 1998). Whether the pre-existing immunological memory at mucosal sites to former instances of respiratory human common cold coronaviruses (e.g. OC43, NL63, HKU1, 229E) is stimulated by intramuscular mRNA boost, panning cross-reactive B-cells remains to be seen (Turner et al., 2021; Wang et al., 2021a).

The approach we introduce here for the evaluation of anti-SARS-CoV-2 humoral response relies on molar units of antigen specific and of total immunoglobulins. Such molar expressions are well-adopted in clinical diagnosis of autoimmune diseases, and provide universal international evaluation and decision making in patients’ care (Monogioudi and Zegers, 2019; Tozzoli et al., 2002). Beyond the obvious benefits of such universality for surveillance and comparative research, our work demonstrates the instrumental importance of absolute units and standardization for mechanistic understanding of functional neutralization. We suggest putting forward this methodological aspect for humoral diagnostics and assessment of vaccine efficiencies in comprehensive universal values.

Comparison of neutralizing activities in serum and saliva upon BNT162b2 vaccination provides the first *in vivo* evidence of augmented specific neutralization of polymeric IgA. At first glance, ‘the multivalent state *per se’* is an obvious explanation following the orthodox proximity-based statistical models of association-dissociation shift – the ‘avidity’ component (Jendroszek and Kjaergaard, 2021). The valence influence is often more prominent for weak affinities, in agreement with the expected shift in the association-dissociation probabilities. Such an avidity component suggests an intriguing possible benefit in the protection toward emerging variants even at the expense of drop in affinities. This classical view is well-studied and has multiple experimental confirmations from comparing kinetic binding and neutralization properties of monovalent Fabs *vs* bivalent IgGs and bivalent IgAs *vs* tetravalent IgAs (Terauchi et al., 2018). Nonetheless, we would like to emphasize that the geometric match between the glycoprotein matrix on the virion surface, and the antibody architecture may significantly impact the neutralization efficiency, given the molecular dimensions of the spike protein. Several recent structural studies have employed cryoelectron tomography to analyse spatial stereochemistry of the authentic SARS-CoV-2 particles (Klein et al., 2020; Yao et al., 2020). Remarkably, the reported center-to-center distances between the trimeric spike foci on the virion surface peak at around 20-30nm, matching the 25nm-longitudinal axis of the dimeric IgA (Figure 5-IIB).

Several recent studies reported the presence of powerful mucosal IgA in post COVID-19 patients (Cervia et al., 2021; Sterlin et al., 2021). Furthermore, Wang *et al* have recently shown that in a recombinant setup, monoclonal IgAs subcloned from circulatory PBMCs of recovered COVID-19 patients exhibit elevated neutralizing potential upon co-expression with the dimer-forming joining chain(Wang et al., 2021b). Recent biotechnological studies have established *ex vivo* systems for the efficient recombinant production of dimeric IgA containing the secretory component.

Whereas the current study illuminates the mucosal aspects of BNT162b2 mRNA vaccine, many additional vaccines are already implemented. While systemic immunity of these vaccines is often thoroughly compared (Collier et al., 2021), their mucosal components are much less explored (Chan et al., 2021). Many more are in clinical trials and development, being of different introduction routes, including mucosal administration, holding a promise of eliciting long-term sterilizing immunity (van Doremalen et al., 2021; Elkashif et al., 2021; Hassan et al., 2021; Lapuente et al., 2021; Tiboni et al., 2021).

In conclusion, our data reveal the existence and the unprecedented specific neutralization potency of spike-targeting temporary mucosal secretory IgA in saliva of BNT162b2 vaccinees. Moreover, our approach of molar quantification of SARS-CoV-2 immunoglobulins in various body fluids may have practical implications for basic research, as well as for accurate assessment of humoral immunity in diagnostics and in epidemiological surveillance studies. Surveying salivary IgA is non-invasive and easily accessible and as such may be beneficial in the search for correlates of protection. Therefore, if predictive, it can be used for large scale, or individual screening, and for determining the need of an additional boost.

## Acknowledgements

We would like to thank for the generous support of the research from The Edmond and Benjamin de Rothschild Foundation. The research was also supported by grants from Israel Science Foundation (338/19), Israel Ministry of Science and Technology (2020000381) and by Science Committee of the Ministry of Education and Science of the Republic of Kazakhstan (AP08856811).

We are grateful for the collaborative effort between different Faculties of the Hebrew University and Hadassah Medical Center for joint research. In particular, we would like to thank the following groups for contributing their time, resources and allocating their members for the research: Prof. Albert Taraboulos, Prof. Sigal Ben-Yehuda, Prof. Ilan Rosenshine, Prof. Emanuel Hanski, Prof. Yuval Dor, Prof. Hanah Margalit, Prof. Ora Furman, Prof. Shoshy Altuvia, Dr. Netanel Tzarum.

We are thankful for help in assay developments, for helpful discussions, ideas, comments on the research and help with writing the manuscript and interpreting the data for: Prof. Albert Taraboulos, Prof. Herve (Hillel) Bercovier, Prof. Sigal Ben-Yehuda, Prof. Ilan Rosenshine, Prof. Zichria Zakay-Rones, Prof. Gilad Bachrach, Dr. Viviana Scaiewicz, Dr. Sharon Karniely, Prof. Moshe Kotler, Prof. Ran Nir-Paz, Dr. Netanel Tzarum, Dr. Michael Berger, Dr. Maayan Salton, Dr. Yoav Shaul, Prof. Mark Saper. We would like to thank Prof. Moshe Kotler, Dr. Nir Paran, Dr. Moshe Dessau and Prof. Amos Panet for the help with reagents and plasmids used for the initial establishment of a variety pseudotype assays. We are thankful to the Hadassah blood bank members and the Hadassah Clinical virology lab members. We would like to especially thank to all our cohort participants and to blood bank donors for their readiness and collaboration.

## Materials and Methods

### Cell lines

Vero E6 and HEK293T cell lines were obtained from the American Type Culture Collection (ATCC). Vero E6 and 293T cells were grown in DMEM medium supplemented with 10% (v/v) Foetal Bovine Serum (FBS), 100 IU/ml of penicillin and 100 mg/ml of streptomycin sulfate, and the cells were grown in 5% CO2 and 95%air. Cells were passaged at 80% confluence and seeded as indicated for the individual assays. Proteins were produced in Expi293 or ExpiCHO that were obtained from Thermo Fisher and grown according to the manufacturer’s instructions.

### Clinical cohort and sample collection

From December, 2020 to May, 2021 we enrolled medical and research personal at our university medical center, without previous documented COVID-19, to participate in our study. Eligible participants were both male and female adults prior to or after receiving the BNT162b2 vaccine. The vaccine has been provided as a part of ongoing national vaccination campaign. This study was part of an ongoing study and was reviewed by the Institutional Review Board (0278-18-HMO). All the participants provided written informed consent. Serum samples were obtained from each participant; saliva samples were obtained from a portion of the participants (saliva cohort). Multiple serum samples were obtained from some of our participants in order to investigate the kinetics of immunoglobulins response to the vaccine (longitudinal cohort). These samples were collected in the following time frames indicated in the cohort tables (supplementary) starting from day 0 till day 180 after the first vaccine. Convalescent plasma and pre-COVID-19 era serum samples were provided from the sample bank and used as a reference for the serum studies; saliva from unvaccinated persons was used as reference for saliva studies. The collected samples were kept at -70⁰C (sera), - 20⁰C (saliva). They were heat inactivated and filtered prior to ELISA or neutralization assays.

### Constructs and plasmids

The following plasmids were used for VSV-pseudovirus production: pVSV-ΔG-GFP, pCAGGS-G or pBS-N-Tϕ, pBS-P-Tϕ, pBS-L-Tϕ and pBS-G(Whitt, 2010).

The plasmids for expression of the Receptor Binding Domain of SARS-CoV2 spike (RBD) and full-length spike (SARS-CoV-2 S (Δ19 aa) were cloned into the pcDNA3.4 backbone (Thermo). The sequences were amplified from SARS Spike synBio (SARS-CoV-2 (2019-nCoV) plasmid using specific primers. The amplified PCR fragments were subsequently cloned using Gibson assembly reaction into pcDNA3.4 backbone modified to include C-terminal Strep Tag-II (IBA). The sequence of the cloned plasmids was verified using Sanger sequencing. All plasmids were amplified under ampicillin selection in Top10 cells (Invitrogen) and purified by NucleoBond Xtra Midi EF kit (MACHEREY-NAGEL).

### Protein production and characterization

Receptor Binding Domain (RBD) of SARS-CoV-2 spike was expressed in mammalian expression system (Expi293 or ExpiCHO) as a secretory protein with sreptag and subsequently purified using streptactin affinity chromatography, followed by preparative size exclusion chromatography on Fast Protein Liquid Chromatography (FPLC) system. The purity and molecular weight of the antigen in solution were verified using SDS-PAGE (Sodium-Dodecyl-Sulfate Polyacryl-Amide Gel-Electrophoresis) and Size-Exclusion-Chromatography Multiple-Angle Laser-Scattering SEC-MALS, respectively.

#### Direct Enzyme-linked immunosorbent assay (ELISA)

Antigen (RBD) was coated onto MAXISORB 96 wells plates in antigen dilution buffer (Tris pH7.5 20mM, 50mM NaCl). Following overnight (ON) incubation at 4⁰C, plates were rinsed 3X with PBS, then blocked with 3% fat UHT milk:PBS (1:1), 30min, RT. Sera samples, and saliva samples (1:1 with PBS) were heat inactivated (60C, 30min) serially diluted in blocking buffer were then added to the wells, incubated for 45min at RT. Wells were rinsed (3X PBS), incubated with secondary Abs coupled to HRP (anti IgA at 1:2500 or IgG at 1:5000) for 30min, and then wells were developed with 3,3′,5,5′-Tetramethylbenzidine (TMB) substrate. The reaction was stopped with 0.4% H_2_SO_4_ and read at 450nm in ELISA reader (Spark, Tecan)

To expand the dynamic range of the assays, the samples (sera or saliva) were always applied as a series of 2- or 3- fold dilutions, and the data was obtained from the dilutions, resided in the linear OD range (“Normalised OD”).

### Sandwich ELISA for total IgG or IgA

MAXISORB 96 wells plates were coated ON with secondary anti-IgG or anti-IgA antibodies, blocked as for direct ELISA and incubated for 30 min with sera and saliva samples, serially diluted in blocking buffer. The wells were then washed (3X PBS), and incubated with secondary Abs coupled to HRP and developed, as described for direct ELISA protocol.

### Quantitative ELISA

Pure commercial antibody was measured via nanodrop, at 280nm, (ThermoFisher) and then diluted serially to final concentrations of 4.4ng/100ul, 1.464ng/100ul, 0.484ng/100ul, and 0.164ng/100ul. ELISA was performed on these 4 dilutions and then a linear graph was made in order to calculate the conversion between the O.D. value of the sample and the corresponding value in ng/100ul. The R^2^ value of this equation was always at least 0.98. This ladder was included in every experiment so that for each sample falling within the linear range of the ladder, the equation could be used to calculate the value in ng/well. Samples with an O.D. value higher than the least diluted commercial sample or lower than the most diluted commercial sample were excluded from the data set. The ng/well value was then multiplied by the dilution factor of that given sample and converted to molar concentration [M] using the molecular weights of 146 kDa for IgG, 150kDa for IgA, and 424 kDa for dimeric IgA.

### Selective IgG and IgA depletion

Streptavidin magnetic beads were washed 4x with PBS and then incubated for 45 minutes with biotinylated capturing antibody diluted 1:10 in PBS. Following incubation, the beads were washed again 4x with PBS before the addition of the sera or saliva samples. The samples were incubated while rotating at 4⁰C for 45 minutes with the beads before being removed. The samples were then checked via ELISA to determine whether complete and selective depletion had been achieved.

### Preparation of pseudotyped VSV and neutralization assay

VSVdeltaG-GFP single round infectious particles were first generated by pseudotyping with VSV-G-envelope expressed *in trans*. P0 generation was produced according to the original Michael Whitt(Whitt, 2010) protocol with minor modification, using co-transfection of the 5 plasmids (pVSV-ΔG-GFP, pBS-N-Tϕ, pBS-P-Tϕ, pBS-L-Tϕ and pBS-G) into HEK293 cells. Infection of the transfected cells with vaccinia T7 polymerase expressing virus was used to drive cytoplasmic T7-driven transcription and mRNA-capping from the plasmids. P1 generation of VSV-G pseudotyped VSVdeltaG-GFP particles was generated by transfection of pCAGGS-VSV(G), followed by infection with P0 particles. P2 generation of the SARS-CoV-2 spike pseudotyped VSVdeltaG-GFP reporter particles was generated by transfection of HEK293 that were subsequently infected with P1 particles.

#### Cell transfection for pseudotyped virus production

HEK293T cells were co-transfected with all 5 plasmids (see above), carrying defective envelope of VSV plasmid (trans form) and GFP reporter plasmid by using Transporter ^TM^5 transfection reagent (PEI, 40,000Da, PolySciences), according to the provider guidelines. Twelve hours post transfection the cells were infected with recombinant Vaccinia T7 polymerase expressing virus(Fuerst et al., 1986) (kindly provided by prof. Moshe Kotler, The Hebrew University of Jerusalem). P0 pseudo-particles progeny was collected at 48 hours post infection: the supernatant was centrifuged (4500xg, 4°C, 30 min) and filtered (0.1 µm, CA filter) to separate VSV pseudoparticles from Vaccinia virus. The complete removal of infectious Vaccinia virus has been confirmed by end-point titration. The filtrates were aliquoted and stored at -80°C. In the second round of infection to generate P2 pseudo-particles, HEK293T cells were transfected with the plasmid encoding SARS-CoV-2 S (Δ19 aa) by using Transporter^TM^ 5 reagent and twelve hours post transfection infected with VSV-ΔG pseudovirus particles. At 48 hours post transfection. the supernatant was collected and concentrated by Lenti-X™ Concentrator following the manufactures protocol (Clontech, CA). The pellet was resuspended in PBS, aliquoted and stored at -80°C. Vero E6 cells were seeded in 96-well plate to get 75-80% confluence and infected with serially diluted SARS-CoV-2 spike (Δ19aa) pseudovirus. The pseudovirus was titrated by counting the green cells 24 hours post infection.

### Neutralization assay

Serum and saliva samples were heat-inactivated (60°C, 30 min) prior to their use in neutralization assays. Saliva samples were pre-diluted 1:1 with PBS before inactivation to avoid coagulation. Next, the samples were diluted in the cell culture growth medium and filtered by using 0.2 μm, cellulose acetate. Vero E6 cells were seeded in 96-well plate and used for neutralization assay at 75-80% confluency. The sera and saliva samples were serially diluted and subsequently incubated with constant amount SARS-CoV-2 spike pseudotyped virus for 1 hr, 37°C. Next, the mixture was transferred to the monolayer and incubated for 16-24hrs. The green fluorescence signal was observed under the microscope at 18-24h.p.i. The reduction in amount of green-fluorescent cells due to neutralization was calculated in percentage of un-inhibited control infection. The images were captured from several fields of each well and the green cells were calculated by using automated image analysis by ImageJ (NIH). The graphs were plotted to get the 50% neutralization titer (NT50), in GraphPad Prism.

## Supplementary Material

### Intramuscular mRNA BNT162b2 vaccine against SARS-CoV-2 induces production of robust neutralizing salivary IgA

#### Assessment of RBD quality and N-linked glycosylation status and validation of custom ELISA

To evaluate the magnitude and the composition of anti-SARS-CoV-2 acquired humoral immunity, we have developed quantitative ELISA using the recombinant RBD derived from SARS-CoV-2 spike protein (Figure S1A, B). Figure S1A shows single band purity of recombinant RBD. Peptide-*N*-Glycosidase F (PNGase F) sensitivity and the predominant Endoglycosidase H (EndoH) resistance of Asparagine-linked (N-linked) glycans of recombinant RBD, points to appropriate post-translational Golgi-derived glycosylation, as it might occur in the settings of natural infection. Evaluation of RBD by size-exclusion chromatography and multiple-angle laser-light scattering (SEC-MALS) shows monodisperse peak with a molecular weight (MW) of 39.2kDa, corresponding to a monomer in solution (Figure S1B). The RBD glycoconjugate analysis demonstrates that approximately 10% of the MW is attributed to N-linked glycans, highlighting their significance in the RBD surface antigenic properties (Figure S1B). The Pearson correlation coefficient of 0.967 between our quantitative test and the ARCHITECT (Abbott, Illinois, U.S.A.) anti-RBD IgG test, confirmed the reliability of our assay, see Figure S1C, for correlation analysis and Table S3, for cohort details.

#### Rationale and details of the anti-SC quantitative ELISA

Basolateral B-cells in the *lamina propria* generate J-joined dimers of IgA that are then luminally delivered by transcytosis to be secreted at mucosal surfaces. The part of the pIg receptor, cleaved during transcytosis (the secretory component, SC), remains permanently associated with the mucosal IgA and is important for its stabilization. To determine the origin of RBD-specific salivary IgA in vaccinees, we asked whether it contains SC. To this end, we employed anti-SC quantitative ELISA measuring total and anti-RBD secretory dimeric IgA (experimental flow is depicted in Figures 4B, C). The molar values were inferred using pure commercially available dimeric IgA as a reference standard (Figure 4C).

---

**Figure S1:**
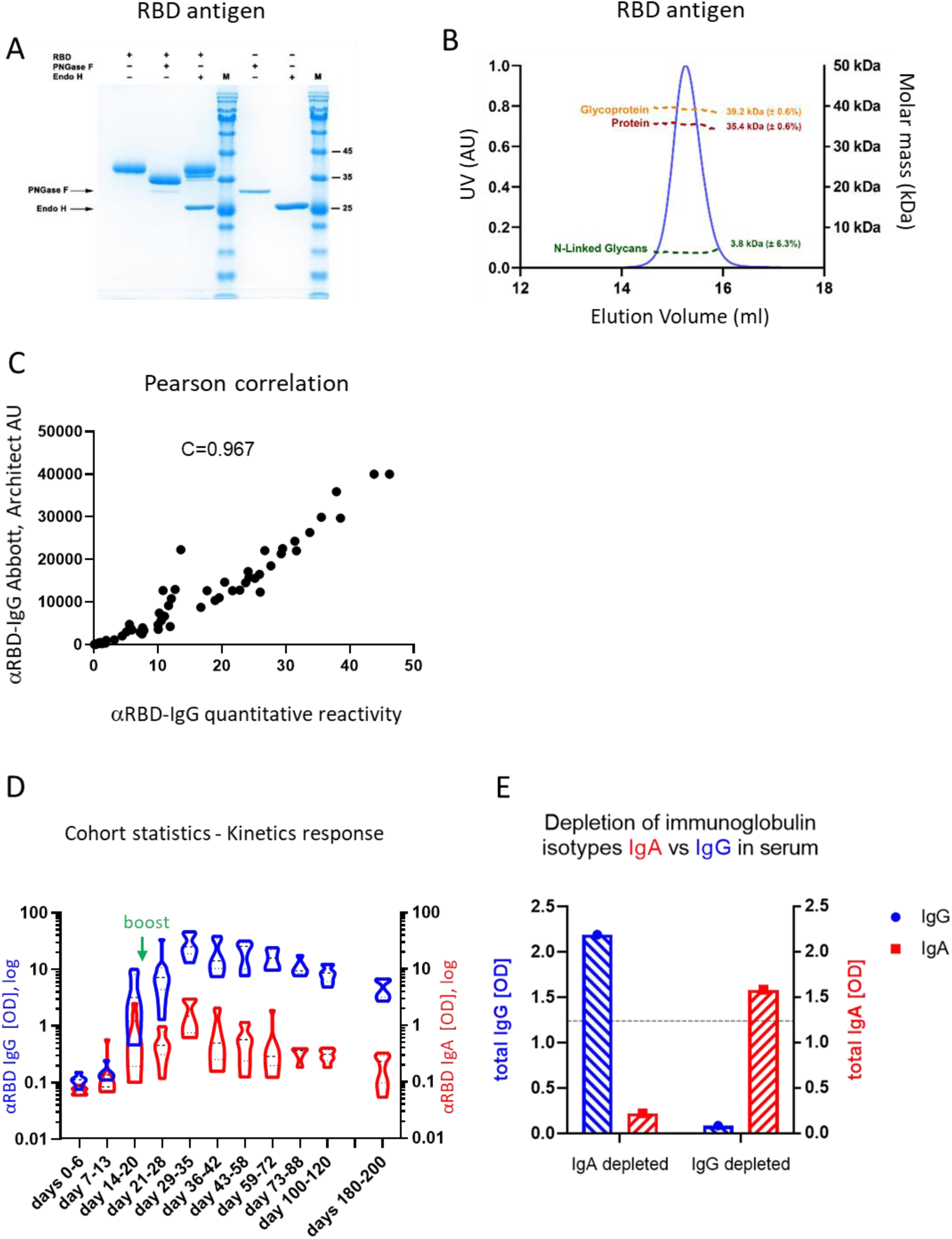
Serological assay: characterization of RBD antigen, validation of anti-RBD ELISA and isotype-specific serological depletion. (A) SDS PAGE of recombinant RBD, produced in mammalian cells in a secreted form. Enzymatic removal of N-linked glycans by PNGase treatment results in characteristic electrophoretic mobility shift, while predominant EndoH resistance demonstrates successful ER-to-Golgi transition during the secretory process. The two lanes on the right show the electrophoretic references of the enzymes (arrows). (B) Size-exclusion chromatography and multiple-angle laser-light scattering analysis confirms monodisperse peak in the form of monomer, with the apparent molecular mass in solution of 39.2kDa, glycoconjugate analysis reveals that the contribution of proteinacious core is 35.4kDa and that the N-linked glycans contribute ∼ 10 percent of the molecular weight of the mature secreted RBD. (C) Pearson coefficient of 0.967 demonstrates linear correlation of our anti-RBD IgG ELISA to routine diagnostic kit of Abbott. (D) Quantitative kinetic profile of anti-RBD IgG (blue) and IgA (red) in serum sampled in the vaccinees cohort, categorized in periods of significance (see Figure 1B, for the uncategorized data presented as a function of days post vaccination). Independent ordinate axes for IgG (left, blue) and IgA (right, red) highlight the restricted, relative nature of the comparison between isotypes in this experiment, as discussed in the text, see also Figure 2 for subsequent developments. Green arrows indicate timing of the second vaccine dose (the boost). (E) Completeness and specificity of IgG depletion from pooled serum samples used for neutralisation experiments presented in Figure 1C. Right-side bars (IgG depleted) serum samples were measured by sandwich ELISA for total IgG (blue column on the right) and IgA (red column on the right), see methods for further details. IgA-depleted samples were measured for total IgG (blue column on the left) and for IgA (red column on the left) were used as a reference for completeness and specificity of IgG depletion. IgG and IgA values are represented on separate ordinate axes (indicated), as explained in the main text. Horizontal dashed line indicates saturation level of ELISA measurement. Total immunoglobulins were measured in this experiment in saturated conditions to confirm the completeness of the depletion.

**Figure S2:**
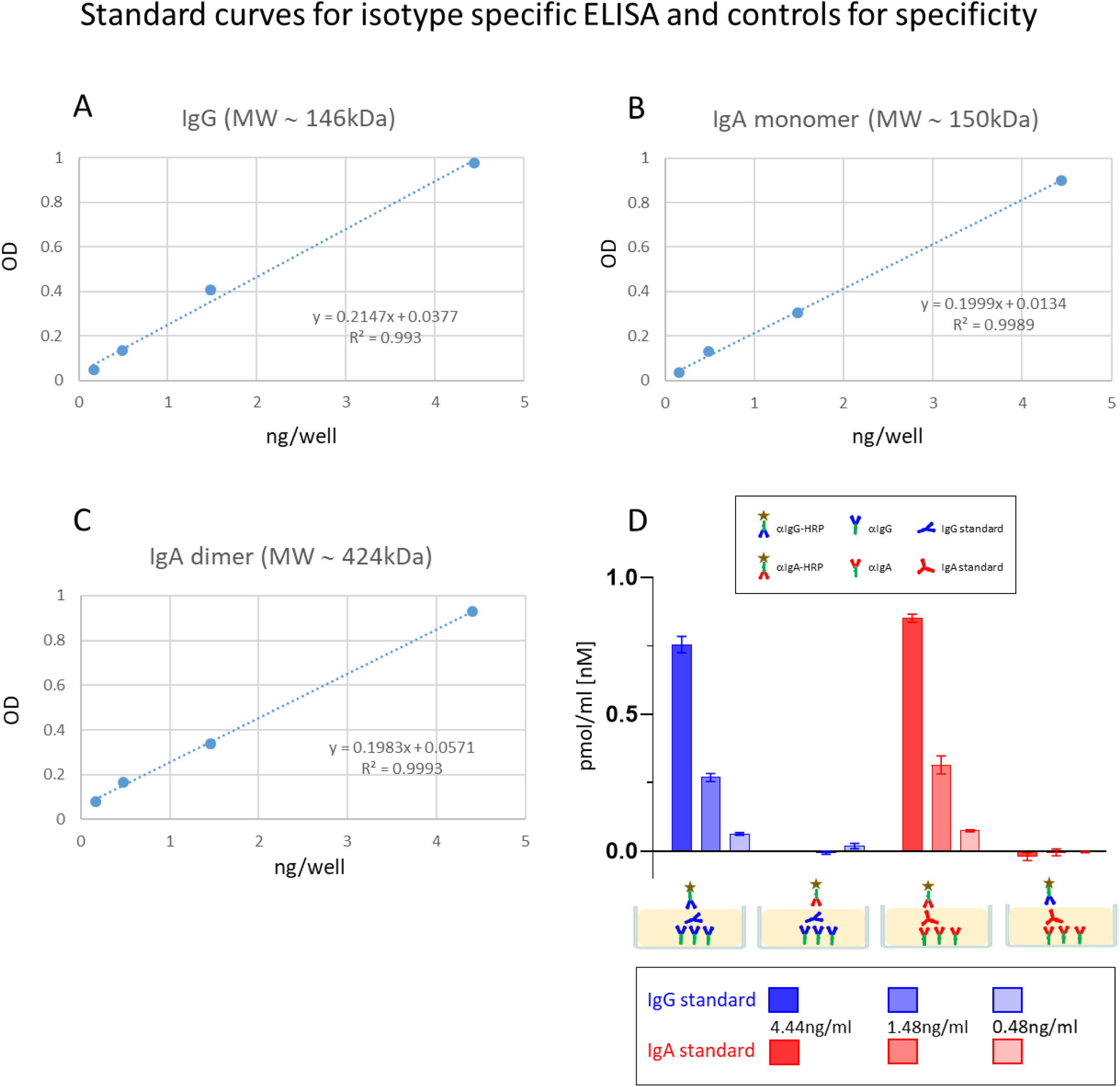
Isotype specific OD-to-mole transformation. (A-C) Sandwich capturing ELISA for selective quantification of total immunoglobulin isotypes (IgG and IgA). Monomeric IgG, IgA and dimeric IgA standards were applied to the plate to transform the OD values to their molar equivalents, shown are standard dilution curves used for extrapolation in capture ELISA format. (D) Shown are control experiments to assess the specificity of isotype capturing and isotype detection antibodies in ELISA assays, as depicted in diagrams above the bars.

**Figures S3:**
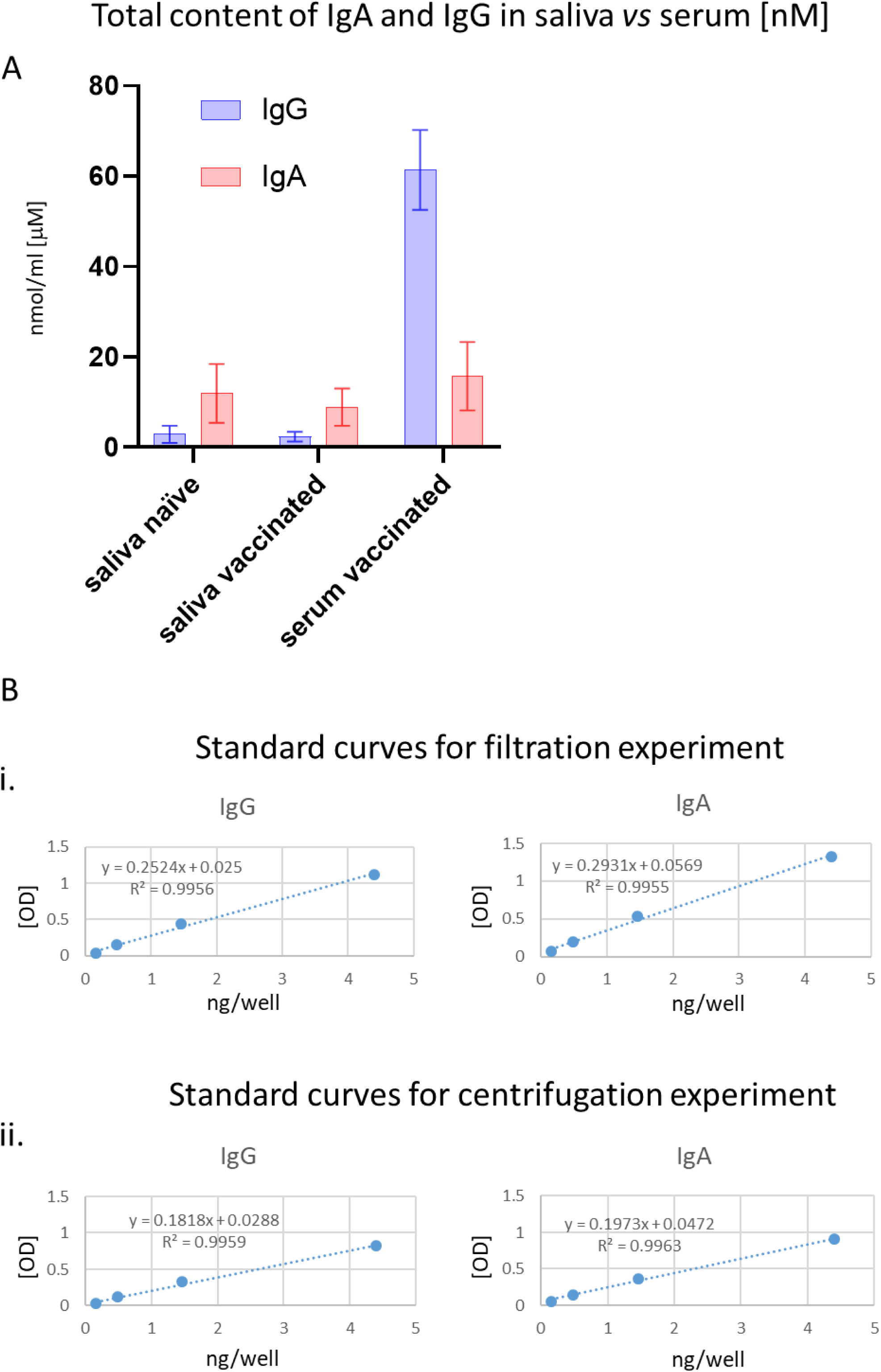
Molar evaluation of the total immunoglobulin content confirms different isotype stoichiometry in saliva *versus* serum. (A) Molar IgA and IgG content were specifically determined in serum and saliva using sandwich ELISA as depicted in Figure 2B (see Methods for details). Tested samples included: naïve saliva (N=13), vaccinated saliva (N=22), vaccinated sera (N=13). IgG is the predominant isotype in serum, while IgA is predominant in saliva in accordance with previously published data. (B) Isotype specific OD-to-mole transformation for saliva samples presented in Figure 3 (see methods for details). Shown are standard dilution curves used for extrapolation in capture ELISA format. Extrapolated molar concentration of used to evaluate recovery yields for soluble anti-RBD IgG and IgA from saliva samples upon (i) centrifugation and (ii) filtration are presented in Table S3.

**Figures S4:**
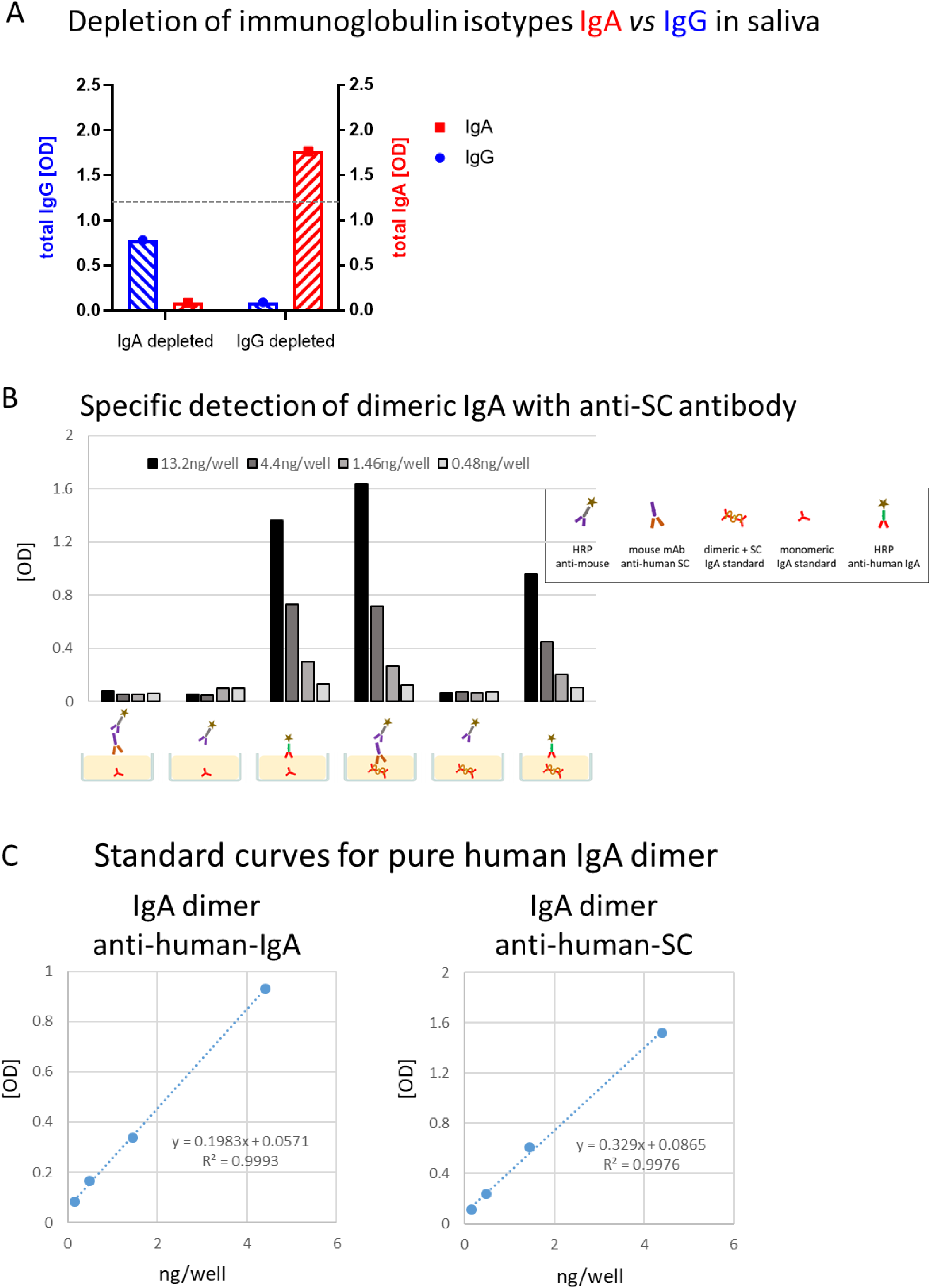
Studies of Abs in saliva of vaccinees. (A) Completeness IgG and IgA selective depletions from pooled saliva samples used for neutralisation experiments presented in Figure 4A. Left-side bars (IgA-depleted) samples were measured by sandwich ELISA for total IgG (blue column on the left) and for IgA (red column on the left). Right-side bars (IgG depleted) samples were measured by sandwich ELISA for total IgG (blue column on the right) and IgA (red column on the right), see methods for further details. IgG and IgA values are represented on separate ordinate axes (indicated), as explained in the main text. Horizontal dashed line indicates saturation level of ELISA measurement. Horizontal dashed line indicates saturation level of ELISA measurement. Total immunoglobulins were measured in this experiment in saturated conditions to confirm the completeness of the depletion. (B) Specific detection of ‘*bona-fide*’ dimeric IgA of mucosal origin bound to Secretory Component (SC) using anti-SC antibody (see methods and diagrams in Figure 4B, C for further details). The experimental details are depicted in diagrams below the corresponding bars. Serial dilutions of commercial standards: (i) monomeric human IgA purified from serum and (ii) dimeric secretory human IgA from colostrum were used as indicated. Graphical legend explains the pictogram identity. (C) Standard curves used for OD to molar transformations in the experiment presented in Figure 4D. Dimeric secretory human IgA from colostrum was used as a standard for detection with anti-human-IgA and anti-human-SC antibodies as indicated.

**Table S1:** **Cohort details related to figure 1A** Serum samples from pre-COVID, vaccinated, and convalescent participants were assayed for anti-RBD IgG and anti-RBD IgA and presented as group in main figure. Here, individual values for each participant are presented. Total number of serum samples (N=90) were taken from 90 participants (P=90) were assayed in the presented sub-cohort. (*) indicates individuals with special immune background. Days pv1 and days pv2 indicate time interval between the respective vaccine dose and the serum sampling. “rec” corresponds to recovered, convalescent, participants. Samples are coded (see more details in the combined cohort table); started at 1-vaccinees and occasional convalescents, starting at 501-convalescent samples cohort; starting from 1001 correspond to pre-COVID cohort.

| Category | Participant | Days pv1 | Days pv2 | Age group | Gender | Anti-RBD I<br>gG (OD) | Anti-RBD<br>IgA (OD) |
| --- | --- | --- | --- | --- | --- | --- | --- |
| preCOVID | 1001 | na | na | 20-40 | f | 0.078 | 0.076 |
| preCOVID | 1002 | na | na | 20-40 | f | 0.085 | 0.077 |
| preCOVID | 1003 | na | na | 20-40 | f | 0.072 | 0.067 |
| preCOVID | 1004 | na | na | 20-40 | f | 0.103 | 0.073 |
| preCOVID | 1005 | na | na | 20-40 | f | 0.090 | 0.067 |
| preCOVID | 1006 | na | na | 20-40 | f | 0.073 | 0.067 |
| preCOVID | 1007 | na | na | 20-40 | f | 0.084 | 0.067 |
| preCOVID | 1008 | na | na | 20-40 | m | 0.084 | 0.068 |
| preCOVID | 1009 | na | na | 20-40 | f | 0.088 | 0.064 |
| preCOVID | 1010 | na | na | 40-50 | f | 0.074 | 0.061 |
| preCOVID | 1011 | na | na | 40-50 | f | 0.112 | 0.070 |
| preCOVID | 1012 | na | na | 20-40 | f | 0.093 | 0.073 |
| preCOVID | 1013 | na | na | 20-40 | f | 0.081 | 0.068 |
| preCOVID | 1014 | na | na | 40-50 | m | 0.088 | 0.068 |
| preCOVID | 1015 | na | na | 20-40 | f | 0.074 | 0.082 |
| preCOVID | 1016 | na | na | 20-40 | f | 0.103 | 0.069 |
| preCOVID | 1017 | na | na | 20-40 | f | 0.078 | 0.062 |
| preCOVID | 1018 | na | na | 60-70 | m | 0.081 | 0.074 |
| preCOVID | 1019 | na | na | 20-40 | f | 0.076 | 0.062 |
| preCOVID | 1020 | na | na | 20-40 | f | 0.090 | 0.065 |
| preCOVID | 1021 | na | na | 20-40 | f | 0.074 | 0.062 |
| preCOVID | 1022 | na | na | 20-40 | f | 0.073 | 0.069 |
| preCOVID | 1023 | na | na | 40-50 | m | 0.083 | 0.068 |
| preCOVID | 1024 | na | na | 20-40 | f | 0.079 | 0.067 |
| preCOVID | 1025 | na | na | 20-40 | f | 0.092 | 0.065 |
| preCOVID | 1026 | na | na | 20-40 | f | 0.082 | 0.062 |
| preCOVID | 1027 | na | na | 20-40 | m | 0.112 | 0.065 |
| preCOVID | 1028 | na | na | - | f | 0.119 | 0.066 |
| preCOVID | 1029 | na | na | 60-70 | m | 0.077 | 0.069 |
| preCOVID | 1030 | na | na | 20-40 | f | 0.106 | 0.068 |
| preCOVID | 1031 | na | na | 40-50 | f | 0.076 | 0.070 |
| preCOVID | 1032 | na | na | 20-40 | f | 0.105 | 0.067 |
| preCOVID | 1033 | na | na | 40-50 | f | 0.078 | 0.065 |
| preCOVID | 1034 | na | na | 40-50 | m | 0.077 | 0.065 |
| preCOVID | 1035 | na | na | 40-50 | m | 0.071 | 0.062 |
| preCOVID | 1036 | na | na | 50-60 | f | 0.089 | 0.066 |
| preCOVID | 1037 | na | na | 60-70 | m | 0.074 | 0.065 |
| preCOVID | 1038 | na | na | 50-60 | f | 0.084 | 0.072 |
| preCOVID | 1039 | na | na | 60-70 | m | 0.077 | 0.066 |
| preCOVID | 1040 | na | na | 40-50 | m | 0.077 | 0.063 |
| preCOVID | 1041 | na | na | 40-50 | m | 0.099 | 0.068 |
| preCOVID | 1042 | na | na | 40-50 | f | 0.100 | 0.073 |
| preCOVID | 1043 | na | na | 40-50 | m | 0.077 | 0.057 |
| preCOVID | 1044 | na | na | - | m | 0.090 | 0.061 |
| preCOVID | 1045 | na | na | 20-40 | m | 0.074 | 0.067 |
| preCOVID | 1046 | na | na | 60-70 | m | 0.078 | 0.073 |
| preCOVID | 1047 | na | na | 40-50 | m | 0.087 | 0.078 |
| preCOVID | 1048 | na | na | - | m | 0.075 | 0.062 |
| preCOVID | 1049 | na | na | - | f | 0.115 | 0.094 |
| preCOVID | 1050 | na | na | 20-40 | f | 0.090 | 0.079 |
| preCOVID | 1051 | na | na | 20-40 | m | 0.095 | 0.067 |

**Table S1: Cohort details related to figure 1A** Serum samples from pre-COVID, vaccinated, and convalescent participants were assayed for anti-RBD IgG and anti-RBD IgA and presented as group in main figure.
| Category | Participant | Days pv1 | Days pv2 | Age group | Gender | Anti-RBD IgG (OD) | Anti-RBD IgA (OD) |
| --- | --- | --- | --- | --- | --- | --- | --- |
| vac | 1 | 32 | 11 | 40-50 | m | 13.192 | 0.609 |
| vac | 2 | 45 | 24 | 40-50 | f | 29.487 | 0.560 |
| vac | 3 | 30 | 9 | 70-75 | m | 25.152 | 2.564 |
| vac | 4 | 51 | 30 | 60-65 | m | 38.551 | 1.719 |
| vac | 5 | 31 | 10 | 50-60 | f | 37.897 | 2.946 |
| vac | 6 | 31 | 10 | 50-60 | f | 18.922 | 2.943 |
| vac | 7 | 40 | 19 | 20-40 | f | 35.532 | 0.658 |
| vac | 8 | 84 | 63 | 40-50 | m | 9.279 | 0.296 |
| vac | 10 | 35 | 14 | 60-65 | f | 43.791 | 0.894 |
| vac | 11 | 35 | 14 | 65-70 | m | 22.785 | 0.744 |
| vac | 12 | 35 | 14 | 20-40 | f | 46.216 | 1.457 |
| vac | 13 | 36 | 13 | 50-60 | f | 11.637 | 0.317 |
| vac | 14* | 42 | 21 | 20-40 | f | 7.524 | 0.154 |
| vac | 15 | 63 | 42 | 65-70 | f | 10.010 | 0.228 |
| vac | 16 | 62 | 41 | 65-70 | m | 13.882 | 0.315 |
| vac | 17 | 63 | 42 | 60-65 | f | 9.707 | 0.121 |
| vac | 18* | 42 | 21 | 60-65 | m | 24.227 | 0.251 |
| rec | 501 | na | na | 40-50 | m | 27.999 | 3.075 |
| rec | 502 | na | na | 20-40 | m | 2.177 | 0.937 |
| rec | 503 | na | na | 40-50 | f | 16.197 | 0.801 |
| rec | 504 | na | na | 20-40 | f | 1.075 | 0.198 |
| rec | 505 | na | na | 20-40 | m | 1.737 | 0.436 |
| rec | 506 | na | na | - | m | 20.085 | 5.497 |
| rec | 507 | na | na | 50-60 | m | 0.285 | 0.083 |
| rec | 508 | na | na | 50-60 | m | 2.160 | 0.170 |
| rec | 509 | na | na | 50-60 | m | 1.853 | 0.259 |
| rec | 510 | na | na | 40-50 | m | 5.148 | 0.308 |
| rec | 511 | na | na | 50-60 | m | 2.721 | 0.324 |
| rec | 512 | na | na | 40-50 | m | 2.535 | 0.090 |
| rec | 513 | na | na | 20-40 | m | 1.363 | 0.234 |
| rec | 514 | na | na | 20-40 | m | 2.464 | 0.172 |
| rec | 515 | na | na | 20-40 | m | 1.199 | 0.185 |
| rec | 516 | na | na | 19 | m | 1.225 | 0.407 |
| rec | 517 | na | na | 20-40 | m | 1.404 | 0.277 |
| rec | 518 | na | na | 40-50 | m | 2.999 | 0.297 |
| rec | 19 | na | na | 20-40 | m | 0.795 | 0.119 |
| rec | 21 | na | na | 20-40 | m | 0.342 | 0.162 |
| rec | 22 | na | na | 50-60 | f | 1.904 | 0.780 |
| rec | 23 | na | na | 20-40 | m | 0.266 | 0.075 |

**Table S2:** **Longitudinal sampling of anti-RBD IgG and IgA in serum - cohort details** Total number of serum samples (N=76 for IgG and N=75 for IgA) in the presented sub-cohort were collected kinetically from 18 participants (P=18). Subjects with serial sampling are indicated in ‘serial sample column’, by consequent numbering. (*) indicates individuals with special immune background. Days pv1 and days pv2 indicate time interval between the respective vaccine dose and the serum sampling. (#) indicates the sample for which only anti-RBD IgG was assayed. See Figures 1B and S2D for the serological anti-RBD IgG and anti-RBD IgA data and the corresponding graphical representations.

| Sample | Serial Sampling | Days pv1 | Days pv2 | Age | Gender |
| --- | --- | --- | --- | --- | --- |
| 1 | I | 0 | na | 40-50 | m |
| 1 | II | 7 | na | 40-50 | m |
| 1 | III | 17 | na | 40-50 | m |
| 1 | IV | 24 | 3 | 40-50 | m |
| 1 | V | 32 | 11 | 40-50 | m |
| 1 | VI | 41 | 20 | 40-50 | m |
| 1 | VII | 48 | 27 | 40-50 | m |
| 1 | VIII | 60 | 39 | 40-50 | m |
| 1 | IX | 84 | 63 | 40-50 | m |
| 1 | X | 108 | 87 | 40-50 | m |
| 1 | XI | 179 | 158 | 40-50 | m |
| 2 | I | 0 | na | 40-50 | f |
| 2 | II | 3 | na | 40-50 | f |
| 2 | III | 7 | na | 40-50 | f |
| 2 | IV | 14 | na | 40-50 | f |
| 2 | V | 21 | na | 40-50 | f |
| 2 | VI | 24 | 3 | 40-50 | f |
| 2 | VII | 28 | 7 | 40-50 | f |
| 2 | VIII | 45 | 24 | 40-50 | f |
| 2 | IX | 56 | 35 | 40-50 | f |
| 2 | X | 84 | 63 | 40-50 | f |
| 2 | XI | 108 | 87 | 40-50 | f |
| 2 | XII | 179 | 158 | 40-50 | f |
| 3 | I | 0 | na | 70-75 | m |
| 3 | II | 14 | na | 70-75 | m |
| 3 | III | 30 | 9 | 70-75 | m |
| 3 | IV | 48 | 27 | 70-75 | m |
| 3 | V | 116 | 95 | 70-75 | m |
| 3 | VI | 186 | 165 | 70-75 | m |
| 4 | I | 0 | na | 60-65 | m |
| 4 | II | 7 | na | 60-65 | m |
| 4 | III | 17 | na | 60-65 | m |
| 4 | IV | 24 | 3 | 60-65 | m |
| 4 | V | 41 | 20 | 60-65 | m |
| 4 | VI | 51 | 30 | 60-65 | m |
| 4 | VII | 56 | 35 | 60-65 | m |
| 4 | VIII | 84 | 63 | 60-65 | m |
| 4 | IX | 108 | 87 | 60-65 | m |
| 4 | X | 179 | 158 | 60-65 | m |
| 5 | I | 7 | na | 50-60 | f |
| 5 | II | 17 | na | 50-60 | f |
| 5 | III | 24 | 3 | 50-60 | f |
| 5 | IV | 31 | 10 | 50-60 | f |
| 5 | V | 41 | 20 | 50-60 | f |
| 5 | VI | 63 | 42 | 50-60 | m |
| 5 | VII | 84 | 63 | 50-60 | f |
| 5 | VIII | 108 | 87 | 50-60 | f |
| 5 | IX | 179 | 158 | 50-60 | f |
| 6 | I | 7 | na | 50-60 | f |
| 6 | II | 17 | na | 50-60 | f |
| 6 | III | 31 | 10 | 50-60 | f |
| 6 | IV | 41 | 20 | 50-60 | f |
| 6 | V | 60 | 39 | 50-60 | f |
| 7 | I | 9 | na | 20-40 | f |
| 7 | II | 20 | na | 20-40 | f |
| 7 | III | 24 | 3 | 20-40 | f |
| 7 | IV | 40 | 19 | 20-40 | f |
| 7 | V | 68 | 47 | 20-40 | f |
| 7 | VI | 103 | 82 | 20-40 | f |
| 7 | VII | 189 | 168 | 20-40 | f |
| 8 | I | 24 | 3 | 40-50 | m |
| 8 | II | 84 | 63 | 40-50 | m |
| 8 | III<sup>#</sup> | 108 | 87 | 40-50 | m |
| 8 | IV | 189 | 168 | 40-50 | m |
| 9 | I | 26 | 5 | 70-75 | m |
| 10 | I | 35 | 14 | 60-65 | f |
| 11 | I | 35 | 14 | 65-70 | m |
| 12 | I | 35 | 14 | 20-40 | f |
| 13 | I | 36 | 13 | 50-60 | f |
| 13 | II | 56 | 33 | 50-60 | f |
| 13 | III | 197 | 174 | 50-60 | f |
| 14* | I | 42 | 21 | 20-40 | f |
| 15 | I | 42 | 21 | 65-70 | f |
| 16 | I | 62 | 41 | 65-70 | m |
| 17 | I | 63 | 42 | 60-65 | f |
| 18* | I | 63 | 42 | 60-65 | m |

**Table S3:** **Pearson correlation cohort details,** including indication of logitudinal sampling, see figure S1C for Pearson correlation between our quantitative test and the ARCHITECT (Abbott, Illinois, U.S.A.) for anti-RBD IgG. Serum samples from naïve, vaccinated, recovered and recovered-vaccinated (vaccinated post-recovery) participants are presented. Total number of serum samples in the presented sub-cohort (N=97) were collected kinetically from 23 participants (P=23). Subjects with serial sampling are indicated in ‘serial sample column’. (*) indicates an individual with special immune background. Days pv1 and days pv2 indicate time interval between the respective vaccine dose and the serum sampling. “rec” corresponds to recovered participants and “rec vac” to participants that were recovered from disease and further vaccinated. Sample numbering is coded and presented according to combined cohort table.

| Category | Participant | Serial sample | Days pv1 | Days pv2 | Age group | Gender | serial sampling in<br>Table S1 for Fig. 1A |
| --- | --- | --- | --- | --- | --- | --- | --- |
| naïve | 1 | a | na | na | 40-50 | m |  |
| naïve | 1 | b | na | na | 40-50 | m |  |
| vac | 1 | c | 0 | na | 40-50 | m | 1 I |
| vac | 1 | d | 7 | na | 40-50 | m | 1 II |
| vac | 1 | e | 17 | na | 40-50 | m | 1 III |
| vac | 1 | f | 24 | 3 | 40-50 | m |  |
| vac | 1 | g | 32 | 11 | 40-50 | m | 1 V |
| vac | 1 | h | 41 | 20 | 40-50 | m | 1 VI |
| vac | 1 | i | 48 | 27 | 40-50 | m | 1 VII |
| vac | 1 | j | 60 | 39 | 40-50 | m | 1 VIII |
| vac | 1 | k | 84 | 63 | 40-50 | m | 1 IX |
| naïve | 2 | a | na | na | 40-50 | f |  |
| naïve | 2 | b | na | na | 40-50 | f |  |
| vac | 2 | c | 0 | na | 40-50 | f | 2 I |
| vac | 2 | d | 3 | na | 40-50 | f | 2 II |
| vac | 2 | e | 7 | na | 40-50 | f | 2 III |
| vac | 2 | f | 14 | na | 40-50 | f | 2 IV |
| vac | 2 | g | 21 | 0 | 40-50 | f | 2 V |
| vac | 2 | h | 24 | 3 | 40-50 | f | 2 VI |
| vac | 2 | i | 28 | 7 | 40-50 | f | 2 VII |
| vac | 2 | j | 45 | 24 | 40-50 | f | 2 VIII |
| vac | 2 | k | 56 | 35 | 40-50 | f | 2 IX |
| vac | 2 | l | 84 | 63 | 40-50 | f | 2 X |
| vac | 3 | a | 0 | na | 70-75 | m | 3 I |
| vac | 3 | b | 14 | na | 70-75 | m | 3 II |
| vac | 3 | c | 30 | 9 | 70-75 | m | 3 III |
| vac | 3 | d | 48 | 27 | 70-75 | m | 3 IV |
| naïve | 4 | a | na | na | 60-65 | m |  |
| vac | 4 | b | 0 | na | 60-65 | m | 4 I |
| vac | 4 | c | 7 | na | 60-65 | m | 4 II |
| vac | 4 | d | 17 | na | 60-65 | m | 4 III |
| vac | 4 | e | 24 | 3 | 60-65 | m | 4 IV |
| vac | 4 | f | 51 | 30 | 60-65 | m | 4 V |
| vac | 4 | g | 51 | 30 | 60-65 | m | 4 VI |
| vac | 4 | h | 56 | 35 | 60-65 | m | 4 VII |
| vac | 4 | i | 84 | 63 | 60-65 | m | 4 VIII |
| naïve | 5 | a | na | na | 50-60 | f |  |
| vac | 5 | b | 0 | na | 50-60 | f |  |
| vac | 5 | c | 7 | na | 50-60 | f | 5 I |
| vac | 5 | d | 17 | na | 50-60 | f | 5 II |
| vac | 5 | e | 24 | 3 | 50-60 | f | 5 III |
| vac | 5 | f | 31 | 10 | 50-60 | f | 5 IV |
| vac | 5 | g | 51 | 30 | 50-60 | f | 5 V |
| vac | 5 | h | 63 | 42 | 50-60 | f | 5 VI |
| vac | 5 | i | 84 | 63 | 50-60 | f | 5 VII |
| naïve | 6 | a | na | na | 50-60 | f |  |
| vac | 6 | b | 7 | na | 50-60 | f | 6 I |
| vac | 6 | c | 17 | na | 50-60 | f | 6 II |
| vac | 6 | d | 31 | 10 | 50-60 | f | 6 III |
| vac | 6 | e | 41 | 20 | 50-60 | f | 6 IV |
| vac | 6 | f | 60 | 39 | 50-60 | f | 6 V |
| naïve | 7 | a | 1- | na | 20-40 | f |  |
| vac | 7 | b | 9 | na | 20-40 | f | 7 I |
| vac | 7 | c | 20 | na | 20-40 | f | 7 II |
| vac | 7 | d | 24 | 3 | 20-40 | f | 7 III |
| vac | 7 | e | 40 | 19 | 20-40 | f | 7 IV |
| naïve | 8 | a | na | na | 40-50 | m |  |
| naïve | 8 | b | na | na | 40-50 | m |  |
| vac | 8 | c | 24 | 3 | 40-50 | m | 8 I |
| vac | 8 | d | 84 | 63 | 40-50 | m | 8 II |
| naïve | 9 | a | na | na | 70-75 | m |  |
| vac | 9 | b | 27 | 6 | 70-75 | m | 9 I |
| vac | 10 | a | 35 | 14 | 60-65 | f | 10 I |
| vac | 11 | a | 35 | 14 | 65-70 | m | 11 I |
| vac | 12 | a | 35 | 14 | 20-40 | f | 12 I |
| naïve | 13 | a | na | na | 50-60 | f |  |
| vac | 13 | c | 36 | 13 | 50-60 | f | 13 I |
| vac | 13 | d | 56 | 33 | 50-60 | f | 13 II |
| vac | 14* | a | 42 | 21 | 20-40 | f | 14 I |
| vac | 15 | a | 63 | 42 | 65-70 | f | 15 I |
| naïve | 16 | a | na | na | 65-70 | m |  |
| vac | 16 | b | 62 | 41 | 65-70 | m | 16 I |
| vac | 18* | a | 42 | 21 | 60-65 | m | 18 I |
| vac | 24 | a | 38 | 17 | 40-50 | m |  |

**Table S3: Pearson correlation cohort details**, including indication of longitudinal sampling, see figure S1C for Pearson correlation between our quantitative test and the ARCHITECT (Abbott, Illinois, U.S.A.) for anti-RBD IgG.
| Category | Participant | Serial sample | Days pv1 | Days pv2 | Age group | Gender | Days between disease onset and blood collection |
| --- | --- | --- | --- | --- | --- | --- | --- |
| rec | 19 | a | na | na | 20-40 | m | 23 |
| rec | 19 | b | na | na | 20-40 | m | 36 |
| rec | 19 | c | na | na | 20-40 | m | 53 |
| rec | 19 | d | na | na | 20-40 | m | 212 |
| rec | 19 | e | na | na | 20-40 | m | 308 |
| rec | 20 | a | na | na | 20-40 | f | 101 |
| naïve | 21 | a | na | na | 20-40 | m |  |
| rec | 21 | b | na | na | 20-40 | m | 27 |
| rec vac | 21 | c | 0 | na | 20-40 | m | 64 |
| rec vac | 21 | d | 12 | na | 20-40 | m | 76 |
| rec vac | 21 | e | 21 | 0 | 20-40 | m | 85 |
| rec vac | 21 | f | 24 | 3 | 20-40 | m | 88 |
| rec vac | 21 | g | 32 | 11 | 20-40 | m | 96 |
| rec vac | 21 | h | 84 | 63 | 20-40 | m | 148 |
| rec | 22 | a | na | na | 50-60 | f | 14 |
| rec | 22 | b | na | na | 50-60 | f | 84 |
| rec vac | 22 | c | 2 | na | 50-60 | f | 101 |
| rec vac | 22 | d | 13 | na | 50-60 | f | 112 |
| rec vac | 22 | e | 22 | 0 | 50-60 | f | 121 |
| rec | 23 | a | na | na | 20-40 | m | 33 |
| rec | 23 | b | na | na | 20-40 | m | 89 |
| rec vac | 23 | c | 14 | na | 20-40 | m | 106 |
| rec vac | 23 | a | 21 | 0 | 20-40 | m | 113 |

**Table S4:**
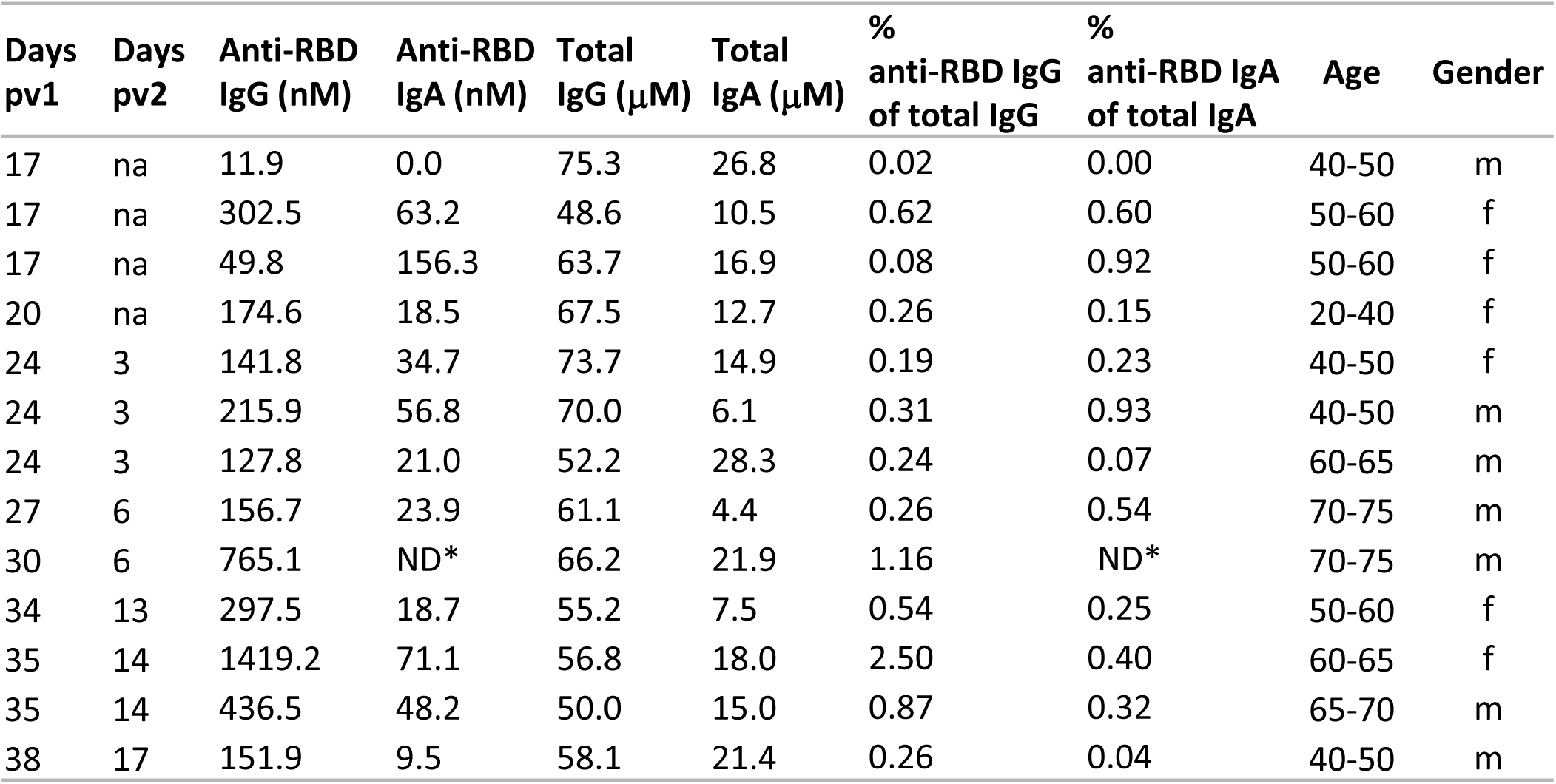
**anti-RBD IgG and IgA vs total IgG and IgA** in serum of BNT162b2 vaccinees expressed in molar concentration (nM and μM, as indicated), see figure 2C and 2D for graphical representation and statistical analysis.

**Table S5:**
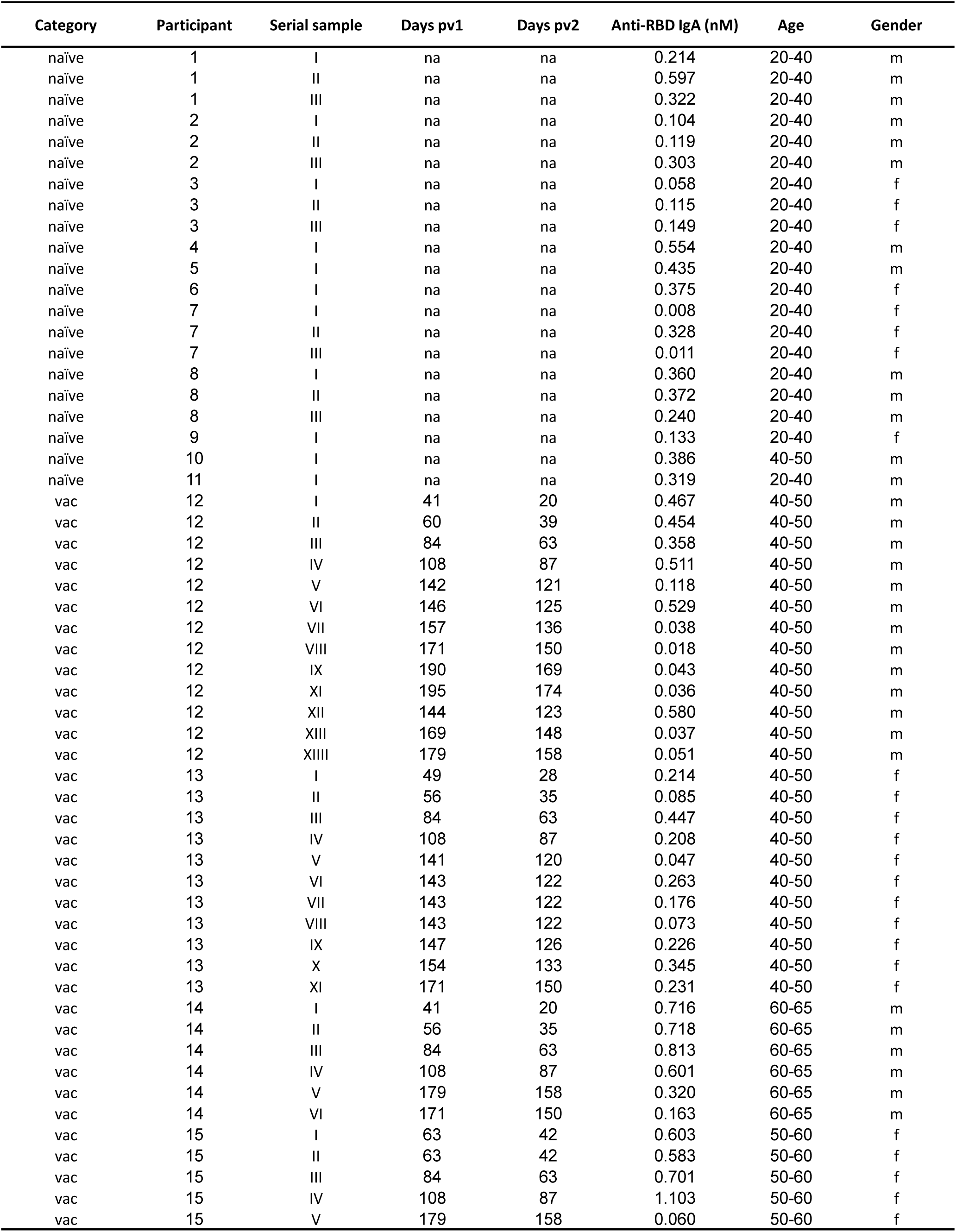
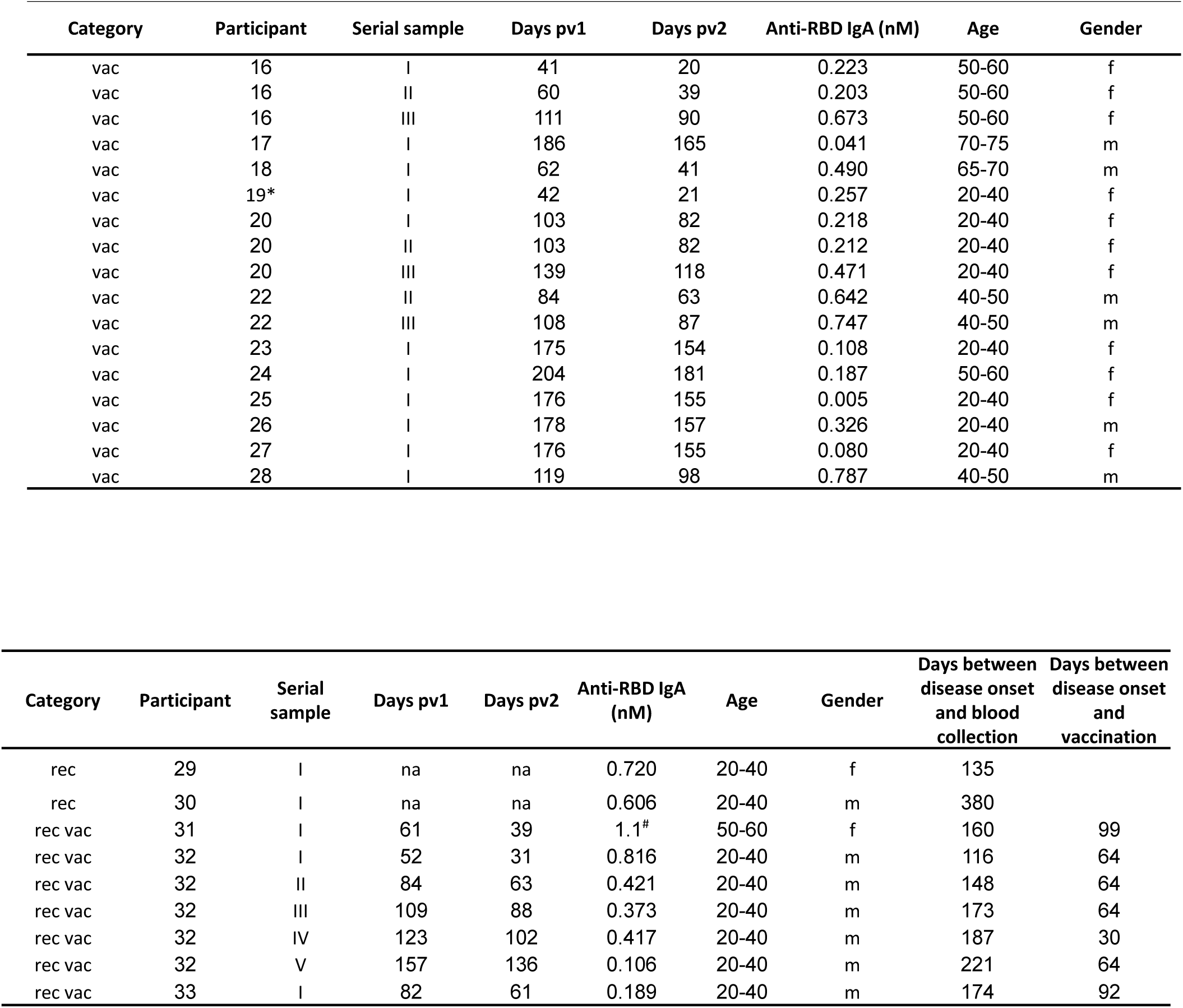
**Saliva cohort details** including indication of longitudinal sampling, see figure 3A for saliva anti-RBD IgA data and the corresponding graphical representations of naïve and vaccinated participants. Total number of saliva samples in the presented sub-cohort (N=82) were collected longitudinally from 33 participants (P=33). **Part I: S**amples from naïve and vaccinated individuals. Samples (N) and participants (P) categorised: unvaccinated (naïve), N(21)/P(11); 40-65d pv1, N(13)/P(8); 80-115d pv1 N(14)/P(8); 135-160d pv1 N(12)/P(3); 170-205d pv1 N(15)/P(10), see Fig.3A in the main text. Subjects with serial sampling are indicated in ‘serial sample column’, by consequent numbering. (*) indicates individual with special immune background. (#) indicates value above the linear detection range. Days pv1 and days pv2 indicate time interval between the respective vaccine dose and saliva sampling. **Part II:** Samples of participants included in the survey that stayed unvaccinated after recovery or recovered and further vaccinated: “rec” corresponds to recovered participants and “rec vac” to participants that were recovered from disease and further vaccinated.

**Table S6:** **Recovery yields of soluble anti-RBD IgG and IgA from saliva samples** tested upon **(i)** centrifugation and **(ii)** filtration. The samples used in panel (ii) were first centrifuged and then either directly assayed or further centrifuged, as indicated.

| Days<br>pv1 | Days<br>pv2 | Anti-RBD IgG (nM)<br>before centrifugation | Anti-RBD IgG (nM)<br>post-centrifugation | Anti-RBD IgA (nM)<br>before centrifugation | Anti-RBD IgA (nM)<br>post-centrifugation |
| --- | --- | --- | --- | --- | --- |
| naive | naive | 0.00 | 0.00 | 0.43 | 0.07 |
| 56 | 35 | 1.41 | 1.68 | 1.56 | 1.38 |
| 108 | 87 | 0.23 | 0.18 | 0.86 | 1.05 |
| 108 | 87 | 0.08 | 0.10 | 0.41 | 0.16 |
| 108 | 87 | 0.39 | 0.31 | 1.56 | 1.50 |
| 123 | 102 | 0.14 | 0.13 | 1.28 | 1.26 |
| 142 | 121 | 0.10 | 0.08 | 0.19 | 0.04 |

| Days<br>Pv1 | Days<br>pv2 | Anti-RBD IgG (nM)<br>before filtration | Anti-RBD IgG (nM)<br>post filtration | Anti-RBD IgA (nM)<br>before filtration | Anti-RBD IgA (nM)<br>post filtration |
| --- | --- | --- | --- | --- | --- |
| naive | naive | 0.00 | 0.00 | 0.17 | 0.24 |
| 56 | 35 | 1.99 | 1.99 | 1.36 | 1.50 |
| 108 | 87 | 0.03 | 0.04 | 0.21 | 0.25 |
| 108 | 87 | 0.46 | 0.52 | 1.67 | 1.48 |
| 123 | 102 | 0.10 | 0.12 | 0.41 | 0.50 |

